# Perceptual detection depends on spike count integration

**DOI:** 10.1101/865410

**Authors:** Jackson J. Cone, Morgan L. Bade, Nicolas Y. Masse, Elizabeth A. Page, David J. Freedman, John H.R. Maunsell

## Abstract

Whenever the retinal image changes some neurons in visual cortex increase their rate of firing, while others decrease their rate of firing. Linking specific sets of neuronal responses with perception and behavior is essential for understanding mechanisms of neural circuit computation. We trained mice to perform visual detection tasks and used optogenetic perturbations to increase or decrease neuronal spiking primary visual cortex (V1). Perceptual reports were always enhanced by increments in V1 spike counts and impaired by decrements, even when increments and decrements were delivered to the same neuronal populations. Moreover, detecting changes in cortical activity depended on spike count integration rather than instantaneous changes in spiking. Recurrent neural networks trained in the task similarly relied on increments in neuronal activity when activity was costly. This work clarifies neuronal decoding strategies employed by cerebral cortex to translate cortical spiking into percepts that can be used to guide behavior.

## Introduction

Elaborating how the brain detects changes in the environment from collections of neuronal responses is essential for understanding neural circuit computations. The onset of even a small, simple visual stimulus drives activity in tens to hundreds of thousands of neurons in sensory cerebral cortex^1^, with some neurons increasing their rate of firing and others decreasing their rate of firing. Increases and decreases in neuronal spiking could both provide equivalent information and be given equal weight by downstream brain regions. However, the brain might have evolved to decode cortical sensory representations using particular response profiles preferentially. For example, the energetic cost of spikes^2, 3^ might favor neural encoding that requires fewer spikes.

That constraint might have driven the development of the visual ON and OFF pathways, where increments and decrements in luminance are converted into parallel channels with low basal rates of firing that respond selectively to changes in only one direction^4^. ON and OFF pathways use fewer spikes to convey the same amount of information compared to an ON-only or OFF-only system^5^. Because cortical neurons similarly have low spontaneous rates of firing, decoding of cortical representations might monitor increases in firing rates preferentially.

In addition to the question of the weight given to different signs of spike rate changes, there is also the question of the relative weight given to rapid versus sustained changes in spike rate. In the periphery, sensory responses often emphasize stimulus onsets and offsets. In space, the edges of stimuli are emphasized by mechanisms like lateral inhibition^6^. In time, the starts and ends of stimuli are emphasized through mechanisms like rapid adaptation^7, 8^. Frequent changes in sensory input like those produced by normal head and eye movements generate transient changes in spiking^9–11^. Decoding of cortical representations might similarly be designed to be sensitive to rapid changes in the rate of firing.

Despite the importance of the question, the relative weight assigned to rapid and slow increments and decrements in cortical spike rates remains poorly understood. Because approaches based on correlations between neuronal activity and behavioral reports cannot provide conclusive results about the weighting of different signals ^12^, this issue requires direct perturbation of neuronal spiking in the cortex of behaving subjects. Optogenetic methods provide an approach for producing controlled increments or decrements in neuronal firing^13, 14^. Studies that have used optogenetics to produce synchronous inhibition of sensory cortex have found that it impairs perception across a range of modalities and stimuli^15–18^. Because electrical excitation of neurons can produce percepts (see^19^), whereas optogenetic inhibition of cortical spiking suppresses perceptual reports, there may be an asymmetry in the ability of increments and decrements of cortical spiking to be used for guiding behavior. However, a direct comparison of the effects of spiking increments and decrements requires that spike rate changes of comparable magnitude be produced and subjects need to be explicitly encouraged to respond to both increments and decrements in signals.

It is critical to understand how downstream neurons readout sensory information from patterns of spiking in upstream populations. Here, we used optogenetic approaches to present increments and decrements in V1 spiking to mice trained to perform visual detection tasks. We report that perceptual detection depends on increases in V1 spiking. When spiking increments and decrements were matched for absolute magnitude in the same neuronal populations, decrements in spiking work against perceptual detection even when they could provide a strong, behaviorally relevant signal. Furthermore, rapid increases in spiking appear to carry no special weight in producing percepts. Instead, perceptual reports appear to depend on the integrated number of spikes rather than instantaneous positive or negative changes in V1 spiking.

## Results

### Using Optogenetic Approaches to Augment Visual Processing During Behavior

In order to compare how V1 spike rate increases and decreases influence perceptual detection, we used optogenetic methods to perturb V1 spiking. Mice were surgically implanted with a headpost and a cranial window to give stable optical access to V1^20^. We used transgenic mouse lines that expressed Cre-recombinase selectively in one of three major subclasses of cortical neurons: excitatory neurons (Emx1, Emx), parvalbumin expressing neurons, PV, and somatostatin-expressing neurons, SST^21–23^. These strains allow selective targeting of excitatory opsins to the neurons of interest with >95% specificity^24, 25^. Following window implantation, we mapped retinotopy in V1 using intrinsic signal imaging (Figure 1A). Imaging data was used to target injections of Cre-dependent viruses containing ChR2-tdTomato^26^ to monocular V1 (Figure 1B,C). Electrophysiological and behavioral experiments were conducted following stable ChR2 expression (≥1 month post-injection).

**Figure 1.**
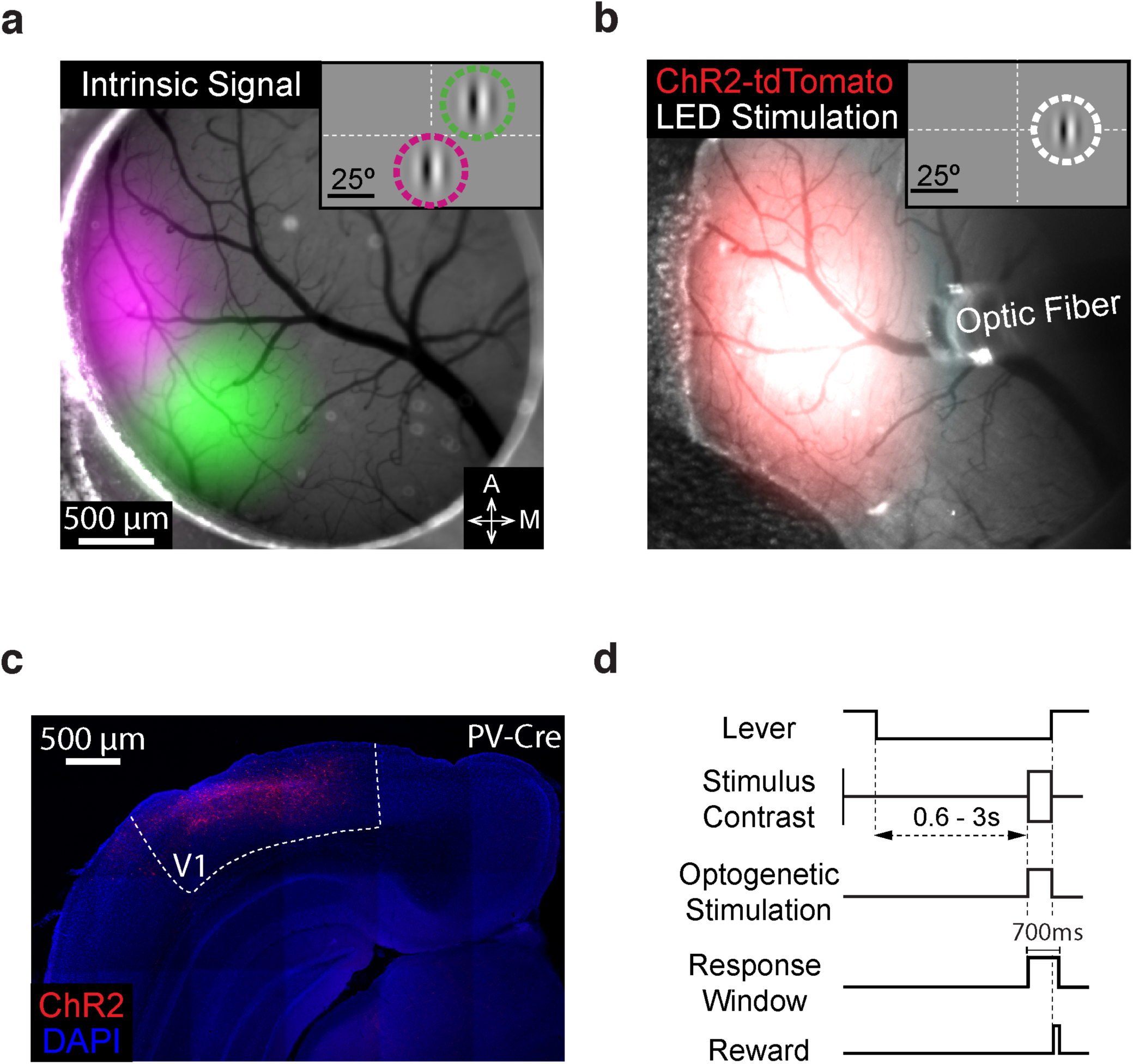
Targeting ChR2 to retinotopically defined areas of visual cortex. A) Pseudo-colored intrinsic autofluorescence responses to visual stimuli presented in two locations in a PV-Cre mouse. Magenta and green features represent 2D-Gaussian fits of responses to stimuli at visual field locations depicted in the inset (magenta: 0° azimuth, −20° elevation; green: 25° azimuth, +20° elevation; Gabor SD = 10°). Dashed lines represent horizontal and vertical meridians. A: anterior; M: medial. B) ChR2-tdTomato fluorescence (2D-Gaussian fit) from the same cortical region shown in A. Area of LED illumination (2D-Gaussian fit) through the optic fiber positioned above ChR2-expressing V1. The retinotopic location corresponding to maximal expression was used in all behavioral sessions (shown in inset; 25° azimuth, 0° elevation; Gabor SD = 6.75°). C) Representative confocal image of ChR2-tdTomato expression in the visual cortex of a different PV-Cre mouse. D). Trial schematic of the contrast change detection task. A single increment and decrement in contrast was selected for stimulation (±15% contrast change for all mice).

In our behavioral experiments, mice were trained to perform a contrast change detection task while head fixed. In this task (Figure 1D), the mouse faced a video display containing static, monochromatic, and vertically oriented 50% contrast Gabor (centered at 20-25° azimuth, −15-15° elevation, 5-7° SD; 0.1 cycles/degree; odd-symmetric) on a mid-level gray background. To start a trial, the mouse depressed and held a lever through a randomly varying delay period (600-3000 ms) after which the contrast of the Gabor increased or decreased (randomly interleaved; 700 ms). The mouse had to release the lever within a 700 ms response window to receive a reward. We randomly varied the magnitude of the contrast change between trials using a range that spanned psychophysical detection threshold. An optical fiber was attached to the headpost to deliver optogenetic stimulation to a consistent cortical location each day (Figure 1B). The optical fiber was aligned with the retinotopic location of the visual stimulus representation in V1 by comparing images of virus expression and intrinsic signal imaging data (Figure 1A,B). Light from the optogenetic stimulus was prevented from cueing the mice to respond using a black fabric shield that attached to the headpost.

### Optogenetic stimulation of principal versus PV neurons produces opposing effects on V1 unit responses and contrast change detection

We first performed electrophysiological recordings from V1 in awake, head-fixed mice (n=8; 4 Emx, 4 PV) to document how V1 units respond to changes in visual contrast and how visual responses were augmented by optogenetic stimulation. Two PV mice were first used in the behavioral experiments that are described later, while the others were prepared only for electrophysiological recordings.

A full-screen 50% contrast sinusoidal grating stimulus (0.1 cycles/degree, static, vertically oriented) was always present on the visual display except during contrast changes. We presented a range of randomly interleaved 500 ms contrast increments and decrements. Spikes were sorted offline and responses to different presentations of each contrast change were averaged. For each stimulus condition, we calculated the change in spike rate (Δspikes/s) by subtracting a pre-stimulus firing rate from the firing rate during a stimulus epoch. When visual stimuli were presented in isolation, firing rate changes were computed using the spike rate from 50-250 ms before stimulus onset compared to 50-250 ms following stimulus onset.

V1 units had diverse responses to contrast changes in the absence of concurrent optogenetic input (n = 250 units; 8 mice). Some units increased their firing rates for either increases (doubling) or decreases (halving) in contrast (Figure S1A; ∼48% of units), while others decreased their firing rates for contrast changes (Figure S1B; ∼33% of units). The mixture of selectivity for increases and decreases in visual contrast is expected given the juxtaposition of ON/OFF receptive fields in V1. Given that we presented gratings of a single orientation and spatial frequency, the visual stimulus was suboptimal for most V1 neurons recorded, but the average population responses to both contrast decrements and increments were decidedly positive changes in firing rate (Figure S1A-D, G-J black traces; Figure S1G). The response was larger for contrast decrements than increments (Figure S1A-D, G-J black traces; Figure S1G). These recordings show that changes in the contrast of a sustained stimulus drive a diverse set of responses in V1 units that could support contrast change detection (Figure S1H), and that V1 is configured so that increases and decreases in stimulus contrast both produce a net increase in spiking (Figure 1; Figure S1G).

On a random half the presentations of 12% (contrast increment) and −12% (contrast decrement) stimuli, contrast changes were paired with concurrent optogenetic stimulation. We used a moderate optogenetic stimulation power (0.3 mW) and visual and optogenetic stimuli were presented for 500 ms. To assess the effects of optogenetic stimulation on visual responses, we calculated change in the average firing rate (Δspikes/s) for each unit using the 50-500 ms before stimulus onset compared to 50-500 ms after stimulus onset. Here, the analysis window was extended relative to above to capture the full duration of optogenetic stimulation. Optogenetic effects were measured by comparing the stimulus evoked firing rates for 12% and −12% contrast changes with and without optogenetic stimulation. To better understand how the effects of optogenetic stimulation compared to the normal physiological response range, we also calculated the change in spike rate for maximal contrast changes (±48% contrast) presented without optogenetic stimulation.

A total of 119 of the 250 units (including multi-units) were recorded across 12 V1 locations in four Emx mice. Large contrast decrements increased overall firing rates (black, Figure 2A,B; -48%: mean +1.6 spikes/s, 0.2 SEM). Moderate contrast decrements drove a weaker response (gray, Figure 2A,B; −12%: mean +0.7 spikes/s, 0.1 SEM). However, when moderate contrast decrements were paired with optogenetic activation of pyramidal neurons, the average response was comparable in magnitude to response for the largest contrast decrement (aqua, Figure 2A,B; −12%+Emx: mean +1.9 spikes/s, 0.2 SEM). Average responses differed significantly between stimulus conditions (Figure 2B; all comparisons at least p < 0.05; Friedman’s test with Dunn– Šidák correction). Thus, optogenetic excitation of V1 Emx-positive neurons significantly increased spiking when paired with moderate contrast decrements.

**Figure 2.**
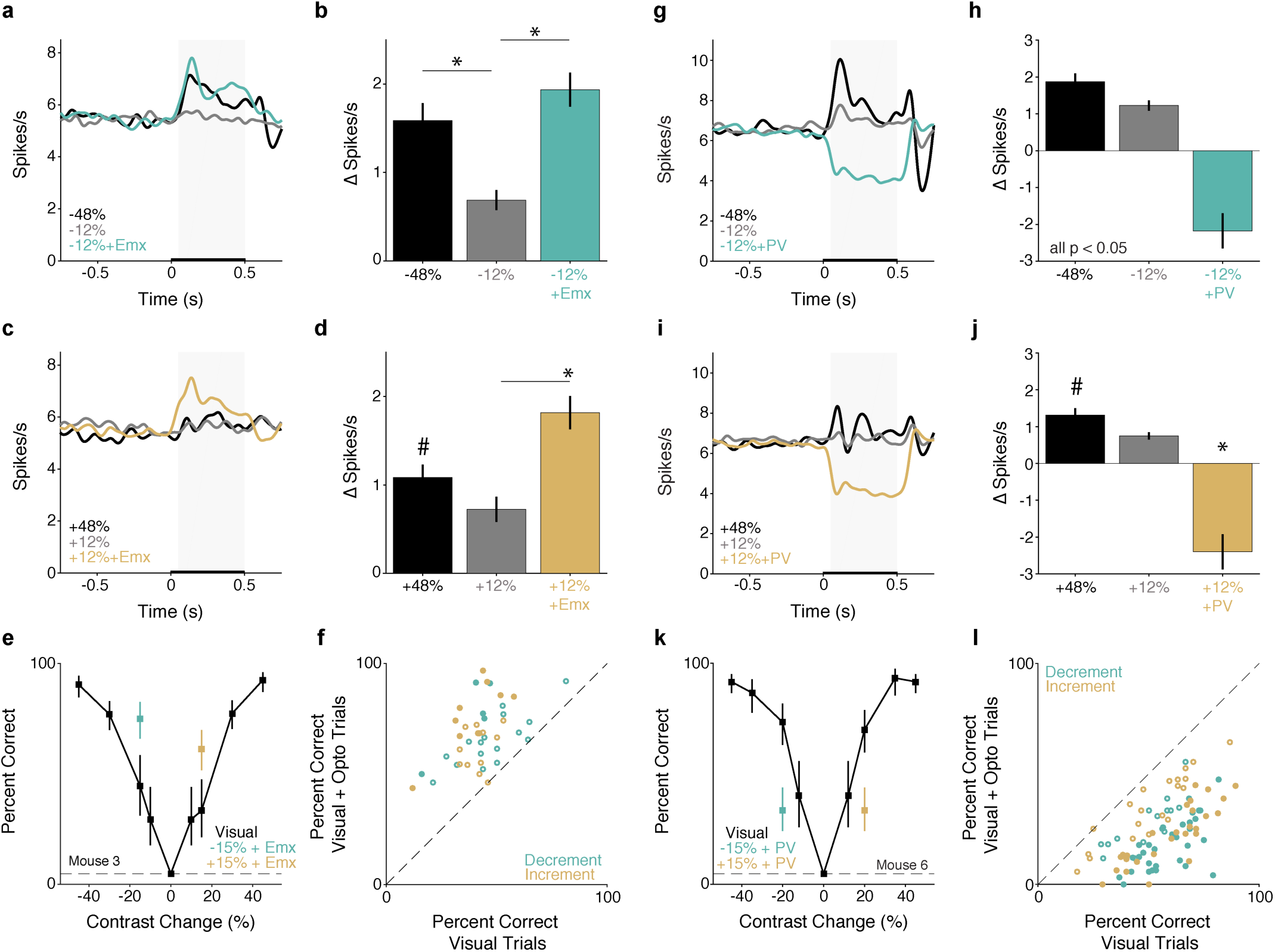
Optogenetic stimulation of principal versus PV neurons produces opposing effects on V1 population responses and contrast change detection. A) Population Gaussian filtered (σ = 25 ms) PSTH in response to large (black) and moderate contrast decrements without (gray) or with (aqua) optogenetic stimulation of pyramidal neurons in Emx mice (n = 119 units). Thickening of the x-axis represents duration of visual and optogenetic stimuli. Gray box = analysis window (+50 -+500 ms) used for spike rate quantification in B. B) Average change in spike rate compared to the time matched baseline period. *p < 0.0001 relative to −12% change; Friedman’s test with Dunn–Šidák correction. C) Same as in A except for contrast increments. D) Quantification of spike rate changes evoked by contrast increments with and without optogenetic stimulation. ^#^p < 0.10 for 48% compared to both 12% contrast change without (gray) and with (gold) optogenetic stimulation. *p < 0.0001 for 12% change with vs. without optogenetic stimulation; Friedman’s test with Dunn–Šidák correction. E) Representative behavioral performance from a single session in an Emx mouse. Points and lines depict the percent correct ± 67% CI for trials without (black) optogenetic stimulation or contrast decrements (aqua) and increments (gold) paired with optogenetic of excitatory neurons. Dashed line = false alarm rate. F) Summary of stimulation effects in Emx mice. Circles depict the percent correct in individual behavioral sessions (3 mice, 21 sessions) with (y-axis) and without (x-axis) optogenetic stimulation, separately for increments (gold) and decrements (aqua). Filled circles = significant change in detection performance (14/42 observations, p < 0.05, Fisher’s exact test). G) Same as in A except for contrast decrements presented with and without optogenetic stimulation of PV interneurons in PV mice (n = 131 units). H) Average change in spike rate with and without stimulation of PV interneurons compared to the time matched baseline period (All comparisons at least p < 0.05; Friedman’s test with Dunn–Šidák correction). I) Same as in G except for contrast increments with and without stimulation of PV interneurons. J) Average change in spike rate evoked by contrast increments with and without optogenetic stimulation of PV interneurons. ^#^p < 0.10 for 48% compared to 12% contrast change without (gray) optogenetic stimulation. *p < 10^-8^ for +12% change with optogenetic stimulation (gold) compared to large (+48%) and moderate (+12%) contrast increments. K) Same as in E, but for a single session in a PV mouse. L) Summary of stimulation effects in PV mice (6 mice, 47 sessions). Conventions are the same as in F. Filled circles = significant change in detection performance (57/92 observations, both increments and decrements; p < 0.05, Fisher’s exact test). See also Table S1, S2, and Figures S1, S2, S3.

V1 responses to contrast increments were weaker than those for decrements (Figure 2C,D, c.f. Fig S1G). Large (48%) contrast increments evoked an overall increase in firing across the V1 population (black, Figure 2C,D; 48%: mean +1.1 spikes/s, 0.2 SEM), while the moderate 12% contrast increment evoked a weaker response (gray, Figure 2C,D; 12%: mean +0.7 spikes/s, 0.1 SEM). As before, when a 12% contrast change was paired with optogenetic activation of pyramidal neurons, the population response was significantly enhanced (gold, Figure 2C,D; 12%+Emx: mean +1.8 spikes/s, 0.2 SEM; p < 0.0001 for 12% change with vs. without optogenetic stimulation; Friedman’s test with Dunn–Šidák correction). Thus, as with contrast decrements, optogenetic excitation of V1 Emx neurons increased the population response to contrast increments.

As expected given the average change in spiking across the population, optogenetic activation of pyramidal neurons significantly changed the firing rate of many units (31%; 37/119, p < 0.05; Wilcoxon’s signed rank test). Of the units significantly modulated by optogenetic stimulation, the vast majority (95%, 35/37) had higher firing rates on trials with optogenetic stimulation compared to trials without stimulation (Figure S1I).

We next asked how optogenetic excitation of V1 during visual stimulation affected detection of contrast changes. Before experimental sessions, all mice were trained to work reliably across a range of interleaved contrast increments and decrements that spanned detection threshold for each change type. During experimental sessions, we delivered optogenetic stimulation on a random half of presentations of a single, near threshold, increment (15%) or decrement (-15%) in contrast. For these measurements, the optogenetic stimulus never exceeded 0.25 mW. The optogenetic stimulus was delayed by 35 ms relative to the onset of the visual stimulus to account for the neuronal response latencies in V1. We restricted optogenetic perturbations to the stimulus epoch so as to only affect V1 spiking responses evoked by the visual stimulus. Both the visual and optogenetic stimulus remained on until the end of the trial so as to prevent stimulus offsets from producing an additional signal that could drive behavioral responses.

Emx mice detected ±15% contrast changes comparably well when those visual stimuli were presented without optogenetic stimulation (median percent correct, decrements 44%, 3% SEM, versus increments 42%, 2% SEM; 21 sessions in 3 mice; p > 0.05, Wilcoxon signed rank test). Optogenetic excitation of pyramidal neurons significantly increased the proportion of trials in which mice detected contrast decrements or increments. Figure 2E shows data from a representative session in which EMX neurons were activated during some trials on which the contrast increased or decreased by 15%. In either case detection was enhanced.

Improvements in detecting contrast increments and decrements were seen in virtually every session (Figure 2F; 21 sessions in 3 mice; decrements median stimulated = 66% [range: 46% – 92%], unstimulated 44% [range: 16%-81%], p < 10^-4^; increments median stimulated = 68% [range: 44% – 97%] versus unstimulated 42% [range: 12%-58%], p < 10^-4^; Wilcoxon signed rank tests). Activation of excitatory neurons significantly increased the proportion of hits in many individual sessions (contrast decrements: 9/21 sessions; contrast increments: 5/21 sessions; p < 0.05, both Fisher’s exact test). Table S1 summarizes session averages for each mouse. Optogenetic activation of excitatory neurons also shortened reaction times on trials in which the animal correctly detected the stimulus compared to trials with no optogenetic stimulation (Figure S2A,B). In summary, optogenetic stimulation of excitatory neurons enhanced the ability of mice to detect changes in contrast regardless of the sign of the contrast change.

The population response to the 12% contrast increments paired with pyramidal neuron stimulation (gold, Figure 2C,D) was larger than the response to 48% contrast increments (black, Figure 2C,D). During behavioral sessions, animals typically detected large contrast increments with greater frequency than trials with optogenetic stimulation (Figure 2E,F). This difference is likely due to our recordings being conducted in different animals outside of the behavioral task, using stimulation powers at the upper limit of those used during behavior.

These observations suggest that perceptual reports for changes in visual contrast preferentially weight increases in V1 spike rates. Previously, we showed that detection of contrast increments is impaired by PV neuron stimulation^16^, but contrast decrements were not examined in that study. The detection of contrast decrements might be mediated, at least in part, by monitoring the spiking of neurons whose firing rates decrease. We sought to test this possibility by reducing V1 spike rates by optogenetically activating PV inhibitory interneurons.

Prior work has shown that activation of PV interneurons in V1 inhibits visually evoked neuronal responses^15, 27, 28^. To confirm that PV-stimulation indeed inhibited population spiking, we recorded from 131 units (including multi-units) from 12 sites in four awake, passively viewing mice with ChR2 expressed in V1 PV interneurons. As with Emx mice, we compared the change in firing rate evoked by large contrast changes (±48%), moderate contrast changes (±12%), and moderate contrast changes with concurrent PV stimulation (±12% + PV; Figure 2G-H). Large contrast decrements evoked a robust increase in firing rate across the population (black, Figure 2G,H; -48%: mean +1.9 spikes/s, 0.2 SEM), whereas moderate decrements evoked a weaker response (gray, Figure 2G,H; −12%: mean +1.2 spikes/s, 0.1 SEM). Pairing moderate decrements with optogenetic activation of PV interneurons robustly decreased V1 output compared to pre-stimulus firing rates (aqua, Figure 2G,H; −12% + PV: mean −2.2 spikes/s, 0.5 SEM). Average responses differed significantly between stimulus conditions (Figure 2H; all comparisons p < 0.05; Friedman’s test with Dunn–Šidák correction). Thus, PV stimulation produced a change in V1 output that was comparable in magnitude, but opposite in sign, to the response to large contrast changes.

Responses to contrast increments were weaker compared to decrements, though still above baseline firing rates. Large contrast increments evoked an increase in V1 output (black, Figure 2I,J; 48%: mean +1.3 spikes/s, 0.2 SEM) and a moderate change evoked a smaller response (gray, Figure 2I,J; 12%: mean +0.7 spikes/s, 0.1 SEM). Pairing a moderate contrast increment with optogenetic activation of PV interneurons strongly and significantly suppressed V1 output compared to pre-stimulus firing rates (gold, Figure 2I,J; 12% + PV: mean −2.4 spikes/s, 0.5 SEM; 12% change with optogenetic stimulation versus other conditions; both p < 10^-8^; Friedman’s test with Dunn–Šidák correction).

Consistent with the strong effect of PV neurons on population responses, most recorded units (65%; 86/131) were significantly modulated by optogenetic stimulation (p < 0.05; Wilcoxon’s signed rank test comparing evoked responses with and without optogenetic stimulation). Of these units, almost all (93%, 80/86) had lower firing rates on trials with optogenetic stimulation compared to trials without stimulation, as expected for activation of inhibitory interneurons (Figure S1J).

To examine how reductions in V1 spiking affected contrast change detection, we prepared and trained a cohort of PV-Cre mice (n=6) to detect interleaved, bidirectional changes in contrast as above. As above, we delivered optogenetic stimulation concurrently with a single, near threshold, increment or decrement in contrast. In the absence of optogenetic stimulation, animals had comparable levels of detection performance for ±15% contrast changes without optogenetic stimulation (median across 47 sessions, decrements 54% correct, 2% SEM versus increments 59%, 3% SEM; 47 sessions in 6 mice; p > 0.05, Wilcoxon signed rank test). Optogenetically stimulating PV interneurons produced behavioral effects that were opposite to those observed in Emx mice. Figure 2K shows data from a representative session in which PV interneurons were activated during some trials on which the contrast increased or decreased by 15%. In either case detection was impaired.

Impairments in detecting increments and decrements were seen in virtually every session (Figure 2L: decrements median stimulated percent correct = 24% [range: 3% – 56%] versus unstimulated 54% [range: 29%-82%], p < 10^-8^; increments median stimulated = 29% [range: 0 – 66.7%] versus unstimulated 59% [range: 18%-90%], p < 10^-8^; both Wilcoxon signed rank test; Figure 2K,L). PV interneuron stimulation significantly decreased the proportion of hits in most individual sessions (decrements: 30/47 sessions; increments: 27/47 sessions; p < 0.05, both Fisher’s exact test; Figure 2L). Table S2 summarizes session averages for each mouse. PV stimulation also affected reaction times, generally slowing responses (Figure S2C,D). These data also argue strongly against the possibility that the facilitation of detection observed in Emx mice was due to the mice seeing scattered light from the optogenetic stimulus.

To further explore these effects, we performed several additional behavioral experiments. As expected based on prior work^15^, the effect of PV stimulation depended on retinotopic alignment between the visual stimulus and optogenetic manipulations. Moving the optical stimulation from the center of the representation of the visual stimulus in V1 reduced the change in performance (Figure S2E-F). Moreover, the effect of stimulation on perceptual reports depended strongly on the power used for stimulation (Figure S2G-K). Furthermore, contrast change detection was also impaired when those changes were presented on the background of a counterphase modulated

Gabor stimulus (Figure S3; Table S3), suggesting that the main effects did not depend on the adaptation of neuronal responses to the static Gabor. Thus, in stark contrast to the results of stimulation in Emx mice, optogenetic stimulation of PV interneurons consistently impairs contrast change perception regardless of sign.

While the effects of optogenetic stimulation through a cranial window are strongest near the cortical surface, we identified ChR2-responsive units throughout cortex (Figure S1K,L) indicating the behavioral consequences of our optogenetic manipulations are unlikely to be restricted exclusively to effects in superficial cortical layers. We also examined how different optogenetic stimulation powers affected V1 spiking and found that the effects of optogenetic stimulation scaled approximately monotonically with power (Figure S1M,N). Together, these data show that our excitatory and inhibitory perturbations on average changes in V1 output that were comparable in magnitude but opposite in sign. Comparing our and electrophysiological and behavioral results strongly suggests that mice preferentially use increments in V1 output to detect changes in visual stimuli.

### Different behavioral consequences of optogenetic increments and decrements of excitatory neuron spiking

The above data show that mice preferentially respond to increments in V1 spiking. However, this may be because spiking increments and decrements were not tested on equal footing. We wondered if mice might exploit spiking decrements if asked to detect changes in sustained, elevated baseline levels of V1 spiking. This approach would also explicitly reward mice for responding to decrements in V1 spiking. Moreover, our prior experiments perturbed different cellular and synaptic elements (excitatory versus inhibitory neurons), which could contribute to the differences in behavioral sensitivity to increments or decrements in overall spiking.

To measure the effects of increments and decrements in optogenetic stimulation on V1 spiking, we recorded from 50 units (including multi-units) from 5 sites in three Emx mice that expressed ChR2. Most V1 units were sensitive to changes in optogenetic input (Figure 3A-C; 62%, 31/50; p < 0.05; Wilcoxon signed-rank on firing rates during the optogenetic step relative to baseline). For the subpopulation of optogenetically-modulated units, firing rates closely followed the optogenetic stimulation (Figure 3A,B). We quantified the evoked change in spike rate during the change in optogenetic input relative to baseline. The firing rate change evoked by decrements and increments in optogenetic input followed the step size (Figure 3C; both p < 10^-4^; Kruskal-Wallis test). Thus, V1 principal neurons can faithfully follow ramps and steps in optogenetic input at modest input powers.

**Figure 3.**
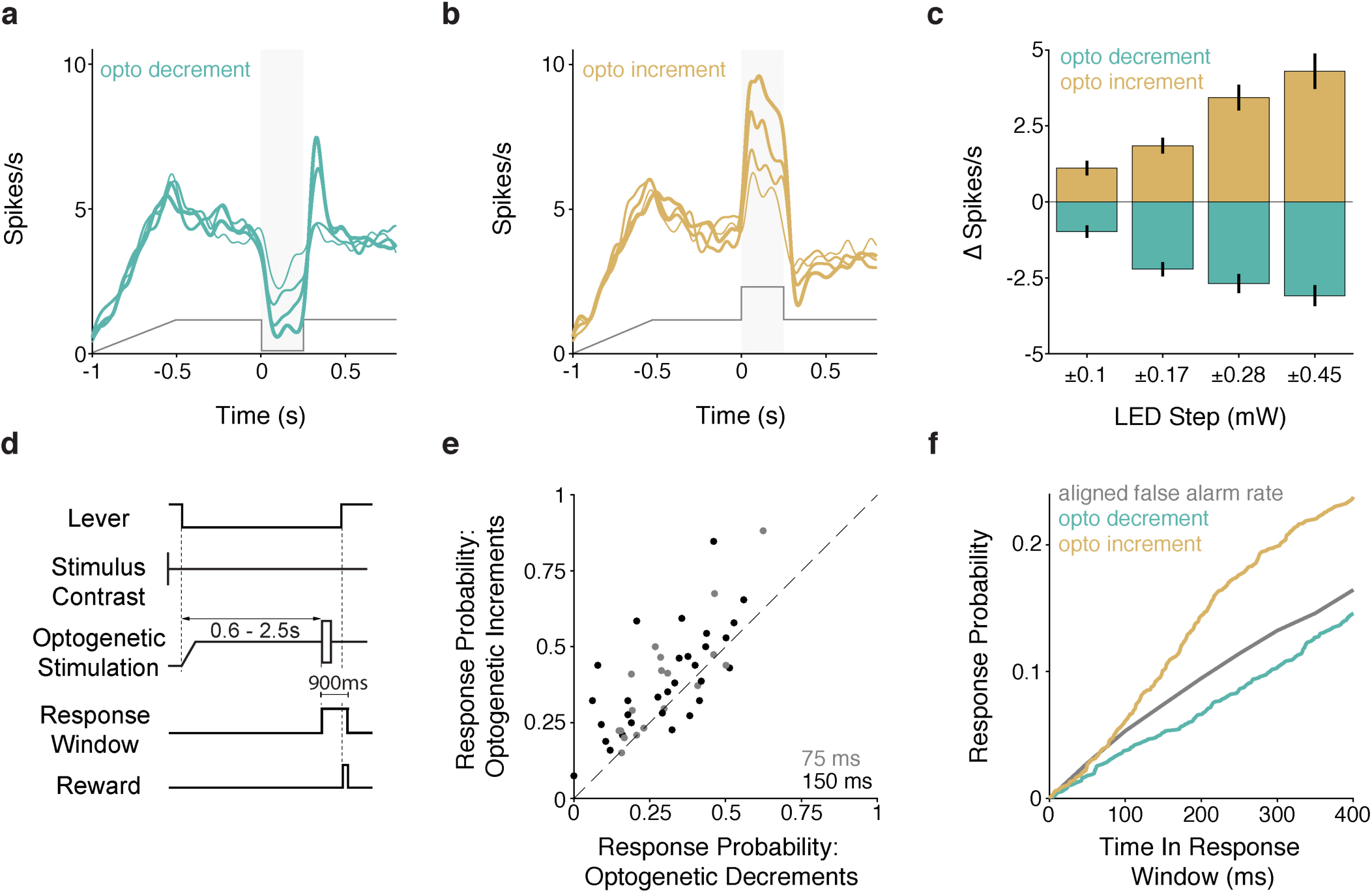
Optogenetically incrementing or decrementing excitatory input into the V1 population asymmetrically affects the probability of behavioral responses. A-C) Electrophysiological recordings confirm V1 spiking follows complex optogenetic input. A) Average PSTHs for optogenetically responsive units (n = 31 units) in response to different magnitude decrements in optogenetic input. Increasing line thickness corresponds to the magnitude of optogenetic decrement (step sizes listed in C). Gray trace depicts the profile of optogenetic stimulation. B) Same as in A, except for increments in optogenetic input. C) Quantification of V1 spike rate changes in response to decrements (aqua) and increments (gold) in optogenetic input. The magnitude of spike rate change significantly depends on the step size (increments and decrements, both p < 10^-4^; Kruskal-Wallis test). D) Trial schematic. The visual contrast was fixed at 50%. At trial onset, the LED power ramped up and then held at a fixed value. After the random delay, the LED power briefly (75 or 150 ms) stepped up or down. E) Scatter plot depicts % correct for trials with optogenetic decrements (x-axis) and increments (y-axis) measured in the same session (n=3 mice; 50 sessions). Gray points = 75 ms optogenetic step duration; Black = 150 ms optogenetic step duration. F) Time course of changes in the lever release probability following optogenetic decrements (aqua) or increments (gold) compared to a time-in-trial matched false alarm rate (gray). See also Figure S4 for data where optogenetic increments and decrements were paired with contrast changes.

We then trained a new cohort of Emx mice on a modified visual detection task (Figure 3D, 3 mice; 50 sessions) that incorporated optogenetic stimulation that persisted through the random delay period. When the visual stimulus changed contrast, the optogenetic stimulus either increased or decreased to potentiate or reduce its input into V1. In addition to examining changes in optogenetic stimulation from a sustained baseline, this design allowed us to examine sensitivity to V1 spiking increments and decrements in the same animals using perturbation of the same circuit (as opposed to effects in different Cre lines).

In all cases, the sustained optogenetic stimulus was relatively modest (0.10-0.25 mW) and stepped up or down by an amount comparable to the baseline power. Our electrophysiological recordings showed that the absolute spike rate changes evoked by decrements and increments within this power range were comparable. The optogenetic stimulus step was brief (75 ms (19 sessions) or 150 ms (31 sessions)), while the contrast change persisted until the end of the trial (900 ms). On a subset of trials, the optogenetic stimulus incremented or decremented after the random delay while the contrast of the visual stimulus did not change. On these trials, the animal was rewarded for responding to increases or decreases in the optogenetic stimulus. Data from trials with concurrent optogenetic and visual stimulation are summarized in Figure S4. In general, mice were better at detecting contrast changes (higher probability of detection, faster reaction times) when the optogenetic input incremented than when it decremented.

We were primarily interested in the behavioral effects on trials where the visual contrast did not change. Mice did not respond reliably to steps in the optogenetic stimulus in the absence of visual stimulus changes, but were more likely to respond to optogenetic increments compared to decrements. Figure 3E shows the session-average probability of lever releases for optogenetic stimulus increases and decreases on trials without visual stimulus changes. The task design encouraged animals to set a liberal response criterion, so many releases were false alarms, but there were far more releases on trials when the optogenetic stimulus increased (increment response rate = 38%, 35-40 95% CI; decrement response rate = 29%, 27-31 95% CI; t-statistic = 8.1, p < 10^-15^, binomial logistic regression). Correspondingly, when mice responded to optogenetic stimuli, reaction times were consistently faster for increments than decrements (Figure S4; increments median: 357 ms, 241 – 585 IQR; decrements median: 467 ms, 257-685 IQR; p < 0.001; Kruskal-Wallis test).

Because the animals were operating with an elevated false alarm rate, we could examine how increases or decreases in the optogenetic stimulus affected the probability of lever releases over time. Figure 3F plots probabilities of lever release following increments (gold) or decrements (aqua) in optogenetic stimulation across all 50 sessions (both 75 ms and 150 ms step durations), together with trial-time-matched false alarm probability (see Methods). The probability of lever responses increased relative to the false alarm rate when the optogenetic stimulus power increased (gold) and decreased when the stimulus power decreased (aqua). This shows that mice are less likely to report a stimulus change when the rate of V1 spiking drops.

Because the optogenetic perturbations were transient, both polarities of optogenetic steps were associated with increases and decreases in spike rate at the step onset (offset) and offset (onset). The firing rate change at the offset of the optogenetic step was sometimes larger than the change at onset owing to overshoot. If perceptual reports were driven by instantaneous increases in firing rate, the end of the optogenetic decrement might be equally well detected as the as the start of the optogenetic increment. However, there was no evidence of a sharp increase in lever releases following the offset of optogenetic decrements (75 – 150 ms in Figure 3F). Instead the lever release probability remained below the false alarm rate for hundreds of milliseconds after the optogenetic decrement despite increases in spike rate that were comparable to those associated with the readily detected optogenetic increments (Figure S4C). This suggests that mice do not respond to instantaneous increases in spike rate relative to immediately preceding epochs of activity, but instead depend on integrating spike counts over periods that likely span hundreds of milliseconds (see Discussion).

### Brief PV or SST interneuron stimulation impairs detection of contrast decrements

The previous experiment suggests that behavioral responses depended on the integrated spike count and not the rate of change in firing rate (derivative of spike rate) in V1. However, the visual stimulus lasted for 900 ms, which could have encouraged mice to adopt an integration-based strategy. Thus, we did an additional experiment in which both visual and optogenetic stimuli were brief (100 ms). Here, we used robust activation of inhibitory interneurons such that the release from inhibition occurred long before the end of the reaction time window. We used higher stimulation powers (0.5 –1.0 mW) to produce a strong reduction in V1 spiking relative to baseline firing (Figure S1M,N). Thus, this stimulation protocol produced a large increase in spiking at the cessation of inhibition.

In addition to changing the temporal profile of inhibition, optogenetic stimulation occurred on 50% of the trials in order to encourage the mice to learn to exploit these brief changes in spike rate in guiding their responses. We also included animals expressing ChR2 in SST neurons to directly compare optogenetic stimulation of different classes of inhibitory interneurons. PV and SST interneurons each suppress V1 responses^28^, but act through distinct synaptic mechanisms^29^. We reasoned that if mice used instantaneous increases in V1 activity to guide their responses, the offset of robust, brief inhibition could potentially facilitate detection. Conversely, if mice rely on the integrated change in spiking, than inhibitory neuron stimulation should always impair detection by subtracting spikes from the population.

For these experiments, we collected new data from PV (n=2; 1 female) and SST (n=2, both male) mice that were part of a previously published report^16^. For the new dataset, mice did a contrast decrement detection task. A 75%-contrast vertically oriented Gabor stimulus (0.1 cycles/deg, centered at 20-25° azimuth, −15-15° elevation, 12° SD, odd-symmetric) was always present on the video display except when its contrast transiently decreased to a lower value (100 ms, Figure 4A). Contrast decrement values spanned psychophysical threshold, and on a randomly selected half of trials, were synchronous with 100 ms illumination of ChR2-expressing PV or SST neurons. We then fit psychometric functions separately to performance values for trials with and without stimulation (see Methods).

**Figure 4.**
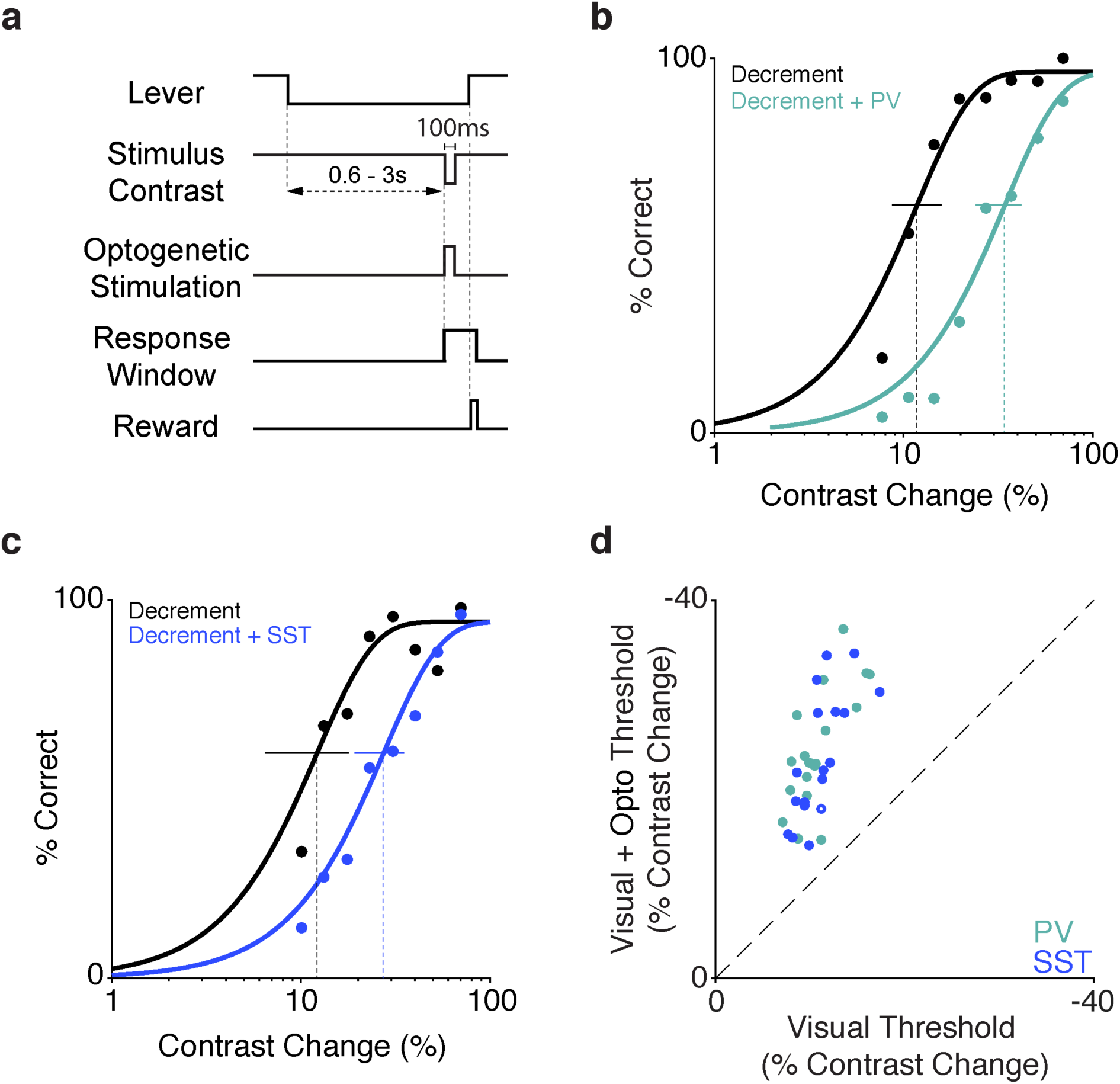
Brief optogenetic stimulation of either PV or SST neurons similarly impairs contrast decrement detection. A) Trial schematic of the contrast decrement detection task. The visual stimulus contrast is fixed at 75% except during a 100 ms decrement. ChR2-expressing interneurons were illuminated with blue light for 100 ms concurrent with the contrast decrement on a randomly selected half of the trials. B) Representative single session performance in the contrast decrement task for a PV mouse. Dots represent false-alarm corrected performance for trials with (teal) and without (black) activation of PV interneurons. Curves are best fitting Weibull functions that were used to determine detection thresholds (dotted vertical lines) and 95% confidence intervals (solid horizontal lines). C) Same as in B but for an SST mouse. Trials with SST stimulation are depicted in blue. D) Summary of PV and SST stimulation effects in the contrast decrement task. Circles depict the contrast decrement detection threshold from individual sessions with (y-axis) and without (x-axis) PV (teal, 2 mice, 20 sessions) or SST (blue, 2 mice, 18 sessions) stimulation. Filled circles represent sessions with a significant shift in threshold (37/38; bootstrapped). See also Figure S2.

Without optogenetic perturbations, there were no detected differences in contrast decrement detection performance between the mouse strains (PV median threshold 11%; SST median threshold 11%; p = 0.89; Wilcoxon rank-sum test). PV or SST activation during contrast decrements impaired detection, shifting the psychometric functions to the right (Figure 4B,C). With the illumination powers used, PV or SST stimulation elevated detection thresholds approximately two-fold (PV 2.4-fold, SEM 0.1, range 1.3–4.2; SST 2.0-fold, SEM 0.1, range 1.3–3.0). This effect is the same as was found previously for contrast increment detection ^16^. Across all sessions (4 mice, 38 sessions), contrast decrement detection thresholds were significantly greater when the visual stimulus was paired with either PV or SST activation (Figure 4D, medians = 23% versus 11%, p < 10^-7^; Wilcoxon signed rank test). This effect was significant for both genotypes individually (PV medians = 23% versus 11%, p < 10^-4^; SST medians = 21% versus 11%, p < 10^-3^; Wilcoxon signed rank tests), and in most individual sessions (PV: 20/20; SST 17/18). Moreover, increasing the stimulation power, which should potentially produce larger instantaneous increments in spiking following the cessation of stimulation, produced larger performance deficits (Figure S2G,H). Combining our previously published data with the current results, PV or SST stimulation never facilitated detection performance across 125 increment or decrement detection sessions (n=8 mice; 5 PV, 3 SST). Not only do these data suggest that mice do not learn to respond to reductions in V1 spike rate, it suggests that any manipulation that removes spikes from the population, even if it includes a response profile with a sharp increase in spiking, will act to suppress the probability of detection.

### Spike costs force recurrent neural networks to preferentially rely on increments in spiking to detect changes in contrast

Preferential readout of spike rate increases compared to decreases might result from constraints that impact the robustness of particular decoding strategies^5^. One potentially strong constraint is the low baseline firing rates of cortical neurons. This limits the dynamic range available for decrements in spike rates to render information to downstream areas and how quickly changes can be detected. As the question of how baseline spiking level influences decoding strategies adopted by neural systems is inaccessible in biological networks, we turned to artificial recurrent neural networks (RNNs).

We trained RNNs (n=50) to perform a contrast change detection task similar to that which we used in mice. RNNs were rewarded for detecting brief decrements and increments in contrast that occurred against an otherwise static baseline contrast (Figure 5A, randomly interleaved). The RNN architecture is described in Methods. Briefly, visual input was passed to a layer of 24 contrast and orientation sensitive units that were either excited or inhibited by visual stimuli (50/50 split). The contrast responsive layer projected to a recurrent layer of 100 units (80 excitatory, 20 inhibitory) and the excitatory units linearly projected onto an output unit that signaled network responses. Because it is likely that the metabolic cost of neuronal activity contributes to the low baseline spike rates in cortical networks^2, 3^, networks were trained using different activity costs. We used rate-based RNNs rather than spiking networks, as recent modeling work suggests that the average firing rate, rather than temporal patterning of spikes, determines the metabolic cost of neuronal activity^30^. This approach allowed us to explore the relationship between activity cost, overall activity, and the decoding strategies used by RNNs to perform the task.

**Figure 5.**
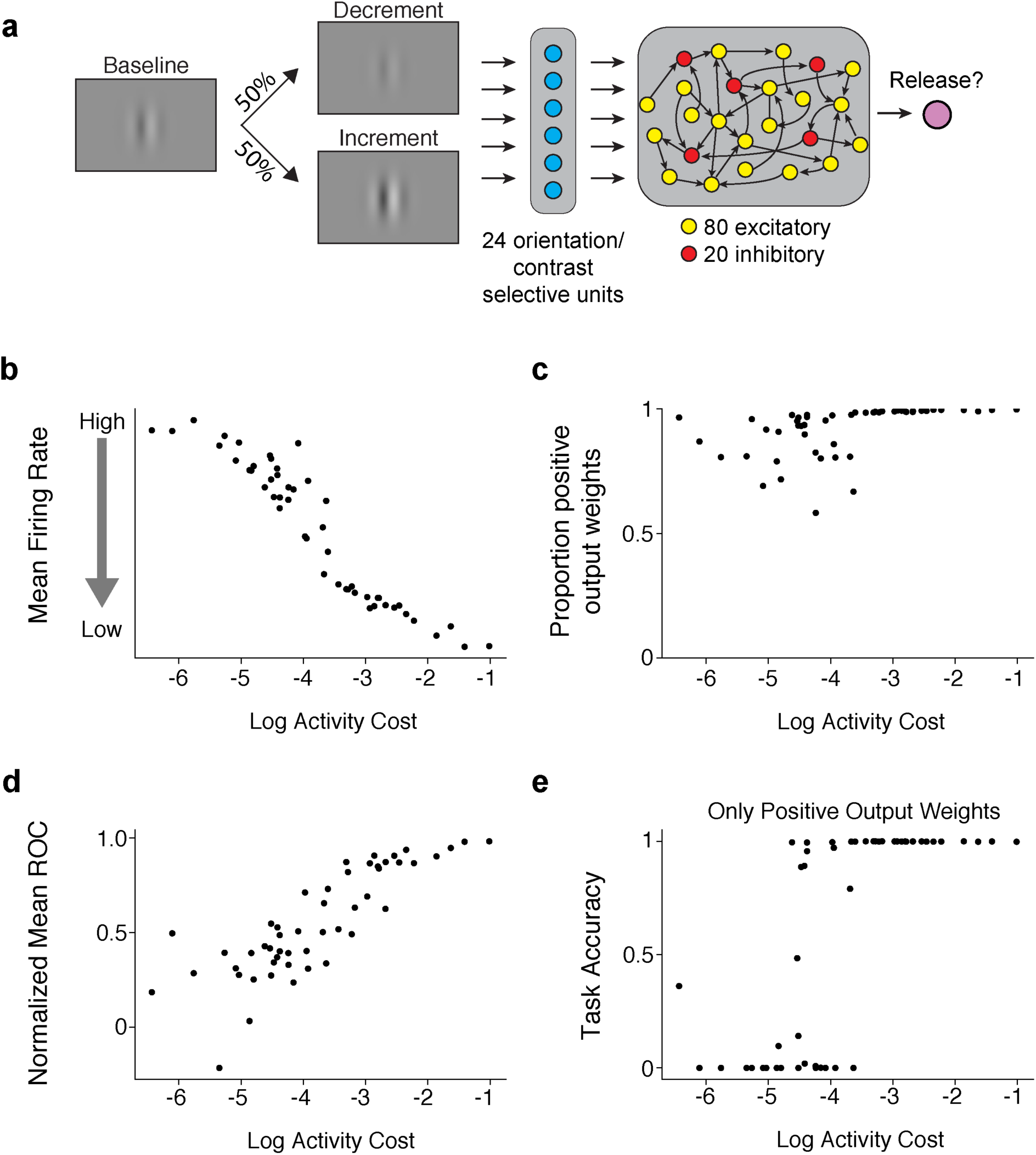
Activity costs force recurrent neural networks to adopt low firing rates and ignore firing rate decrements. A) Schematic of a network trained to detect interleaved increments and decrements in contrast. The input is a single oriented Gabor that increases or decreases in contrast, which is passed to 24 units that are tuned to orientation and contrast (12 excited or inhibited by contrast changes). The contrast responsive layer passes input to a recurrent layer with 100 interconnected units (80 excitatory/20 inhibitory) that project to a single output node that controls lever releases. B) Increasing activity costs (x-axis) drive recurrent networks to lower basal rates of firing (y-axis). Points represent the results obtained from different networks trained with different activity costs. C) The proportion of output weights from the recurrent layer that are positive increases as function of activity cost. D) Normalized ROC (mean of increments and decrements). Range extends from −1 (all units decrease in firing) to 1 (all units increase). E) Task Accuracy as a function of activity cost when all units with negative weights are removed. See also Figure S5.

Activity costs profoundly affected the average firing rate observed in RNNs. As activity costs increased, networks adopted significantly lower average rates of firing (Figure 5B; Spearman ρ = -0.96, p < 10^-27^). Consistent with the idea that lower levels of activity impact the dynamic range available for decrements in spiking to signal stimulus changes, the proportion of positive output weights from the recurrent layer depended strongly on activity cost. High activity costs reliably caused RNNs to use exclusively positive weights (Figure 5C; Spearman ρ = 0.79, p = 10^-10^). This demonstrates that the metabolic cost of activity can reduce firing rates and shift networks toward using firing rate increments.

We next used receiver-operating-characteristic (ROC) measures (see Methods) to explore how changes in RNN activity related to behavioral responses. For each network, we calculated a population ROC based on responses to contrast changes. A value of −1 indicates that all units in a network fired less when RNNs correctly detected contrast changes (hit) compared to trials in which the network failed not respond (miss). Conversely, a value of 1 indicates that activity was higher for all units in the network when the network responded. The normalized ROC value converged to 1 as activity costs increased (Figure 5D; Spearman ρ = 0.85, p < 10^-14^). This shows that behavioral responses to increments and decrements in contrast were correlated with firing rate increases for high activity costs in our RNNs.

To examine the causal role for increases and decreases in activity rate in guiding responses, we presented new trials to trained networks but turned off outputs from the recurrent layer that were either positive or negative. To align with our neurophysiological data in V1, this meant that decreases in spiking still exist and can influence activity within the recurrent layer, but only positive or negative outputs from the recurrent layer to the response neuron could impact its responses. Removing negative outputs devastated performance when activity costs were zero, but removing contribution of negative output weights had almost no impact on performance once activity costs became appreciable (Figure 5E; Spearman ρ = 0.77, p < 10^-10^). Networks had modest performance when activity costs were low and they were constrained to use positive output weights (Figure 5E). However, networks constrained to negative weights performed poorly, even with low activity costs (Figure S5). These data show that activity costs in artificial neural networks can produce decoding strategies that are consistent with those observed in the mouse: Increments in neuronal activity appear to be preferentially used for detecting changes in visual stimuli.

## Discussion

How do downstream brain regions decode changes in cortical spiking? Our data suggest that although decreases in the output of V1 neurons convey behaviorally relevant information, they are not used in supporting the detection of visual stimuli. Subpopulations of V1 units have firing rates that either increase or decrease in response to contrast changes, likely owing to the ON/OFF juxtaposition of early stage visual receptive fields. This demonstrates that either increases or decreases (or both) in neuronal firing could inform decisions about visual stimuli (Figure S1). Optogenetic excitation of primary neurons facilitated contrast change detection for both increases and decreases in contrast whereas optogenetic manipulations that reduce the spiking of V1 neurons impaired detection of contrast changes of either sign (Figures 2-4, S1-3).

The perceptual impairment produced by optogenetic inhibition persisted throughout testing, suggesting that downstream structures cannot readily learn to decode decreases in V1 spiking. The brain might not process reductions in cortical spiking because low baseline firing rates in cortex impacts the coding range for decrements. Indeed, we found that imposing a simple metabolic constraint made RNNs adopt low baseline rates of firing and encouraged decoding strategies based exclusively on increases in activity (Figures 5, S5).

When mice were trained to respond to brief optogenetically-induced increments and decrements of V1 spike rates in the same neuronal populations, only steps that increased the rate of firing above baseline were detected, even though steps of either sign results in rapid increases and decreases in the rate of firing (Figures 3,4). This suggests that detection depends on integrating spike counts over periods of hundreds of milliseconds, and that the derivative of spike rate over much shorter periods cannot be used, regardless of the sign of that derivative (Figures 3,4, S2,3). In sum, our results suggest that downstream circuits detect changes in visual stimuli by integrating spikes from V1 to some criterion value, as has been suggested by the popular drift-diffusion model ^31, 32^.

### Why would cortex preferentially decode from increments in neuronal spiking?

Excitation and inhibition are tightly coupled in cortical circuits^33–36^. Strong inhibition keeps baseline firing rates low, which limits the dynamic range available for decreases in firing rate to encode information. In the visual system, natural scenes generate sparse firing rates in V1^11, 37^. Consequently, normal operating regimes provide only a modest pedestal of spiking in which to decrement spike rates, while enhancing the encoding potential for spike rate increments. These features might be adaptations to the metabolic cost of spiking, as energy usage by the brain depends primarily on the rate of neuronal spiking^2, 3^. Such constraints likely favor energy efficient neuronal codes^5, 38^. Our modeling experiments support this general idea, as increasing the cost of neuronal activity drove RNNs to adopt lower baseline rates of firing and encouraged preferential reliance on increments in neuronal activity to detect changes in visual stimuli (Figures 5, S5).

In the retina, bipolar cells are functionally specialized to segregate increases and decreases luminance into distinct ON and OFF processing streams^4^. This split allows a given number of retinal ganglion cells (RGCs) to convey information more efficiently than a single channel, especially when the costs of spiking are considered^5^. The fact that pharmacological inhibition of ON cells disrupts perceptual reports for light increments without affecting decrements^39, 40^ supports the idea that central structures decode primarily spike increments. Recently, Smeds and colleagues^41^ used a transgenic mouse with differentially elevated luminance threshold of the ON and OFF RGCs to show that mice doing an absolute luminance detection task depend the ON pathway, even when the OFF pathway could provide greater sensitivity. These results strongly support the idea that decoding by central visual structures depends primarily on increases in spiking.

### Implications for signal readout in V1

Primates can learn to report electrical microstimulation of all cortical areas tested^42–46^, but detection improved markedly with extended practice^47^. If mice could decode decreases in V1 output, the effects of PV or SST stimulation might emerge only slowly as mice learn to use the resultant signal to guide responses. However, we previously showed that PV or SST activation consistently elevated contrast increment detection thresholds over many weeks of testing^16^. Here, we extended the training of a subset of these same animals to specifically encourage them to detect spike rate decrements and found that PV or SST activation still produced a perceptual impairment (Figure 4).

These experiments spanned weeks (max 21 sessions, ∼4500 stimulation trials). Other PV mice were tested with different optogenetic stimulus powers or stimulus configurations and testing spanned 20-30 (max = 32) sessions, yet we never observed an increase in detection probability associated with optogenetic activation of interneurons that reduce cortical spiking. For comparison, mice learn to report optogenetic activation of vasoactive intestinal peptide (VIP) expressing interneurons, which potentiate cortical spiking^48^, with far less total exposure^16^ (9-15 sessions; ∼1800-3000 stimulation trials). Moreover, mice can perceive optogenetic activation of V1 excitatory neurons and the change in spike rate produced near the detection threshold is low^49^ (Δ1.1 spikes/unit). Our PV manipulations produced spike rate changes that were similar in magnitude but opposite in sign (Figures 2, S1, S3). Thus, despite thousands of exposures to the population level consequences of PV or SST stimulation, mice do not learn to use this signal to guide their responses.

Brief decrements in excitatory input similarly suppressed behavioral responses, even though those decrements were followed by sharp increases in spiking (Figure 3). These mice were trained using sustained visual stimuli, which may have encouraged them to integrate spikes over long time windows. However, we encouraged a separate group of mice to use brief epochs in spiking to guide their responses. Here, strong suppression of V1 spiking with PV stimulation followed by release from inhibition consistently reduced the probability of behavioral responses (Figure 4). Together, these data argue that mice do not decode instantaneous changes in cortical spiking, but instead guide their behavioral responses by integrating V1 spiking over hundreds of milliseconds. Our observations are consistent with previous work where detection of optogenetic stimulation in V1 was dependent not on the precise timing of inputs but on the total input delivered over short periods (100 ms, Histed and Maunsell, 2014). Thus, even though sensory neurons convert both the spatial and temporal edges of visual stimuli into robust, transient changes in spiking, perceptual reports appear to rely on spike count integration. While accumulation of information closely describes links between neuronal spiking and perceptual decisions in higher order brain areas ^51^, similar mechanisms may underlie the production of percepts throughout sensory cortex, and possibly beyond. Our data also raise important questions is about the timescale of integration and whether it can be flexibly adjusted. Extrapolating from studies of attention and decision-making, it is likely that windows for cortical integration are flexible^52, 53^. Future work should address the periods over which perceptual integration takes place and the extent of its flexibility.

### Interactions Between Visual and Optogenetic Input

How do optogenetic perturbations interact with visual signals in a V1 neuronal population? We previously showed that the effects of PV or SST stimulation on contrast detection were better explained by a divisive scaling of the stimulus contrast compared to models in which stimulation acted directly on the probability of lever releases^16^. This argues strongly against the possibility that interneuron activation impairs performance by disrupting motor planning or produces a signal that distracts the animal. Optogenetic manipulations that increase V1 output might correspondingly increase the apparent contrast of visual stimuli. However, activation of excitatory neurons or VIP interneurons can produce percepts in the absence of any natural visual stimulus^16, 50^. Thus, optogenetic activation of excitatory neurons when a visual stimulus is present could either enhance stimulus representations, or could produce a signal that is distinct from the visual signal. Here, we delivered moderate optogenetic excitation alongside weak (or non-existent) visual stimuli and produced sub-saturating enhancements in detection performance, suggesting that the optogenetic stimulus was unlikely to be readily perceptible in isolation. Work from others suggests that artificial and natural signals are merged into a common percept (see^19^). For example, when electrical microstimulation is delivered to direction selective neurons in the middle temporal area (MT) while monkeys view motion stimuli, the animals’ reports resemble a vector average of the motion stimulus and the preferred direction of the stimulated column^54^. Regardless of the exact mechanism by which potentiating excitation facilitates performance in our experiments, the conclusion is unchanged: adding signal that the brain can decode can increase the probability of detection.

### Implications For Future Work

It remains to be determined how general these results are for cerebral cortex and other brain structures. Increases and decreases in spike output are likely to be equally important for signaling in some other brain areas. In the cerebellum and the basal ganglia aspects of eye position^55^, head position^56^, speed and rotation^57^, and action initiation^58^ all appear to critically rely on decrements in spike rate. Many of these systems exhibit relatively high baseline rates of firing compared to cerebral cortex or signal using inhibitory rather than excitatory neurotransmission. It would be unsurprising if different constraints have shaped diverse sets of information processing strategies across different brain regions.

Determining how changes in spike rate mediate the functions of neural circuits is critical for understanding the brain. Currently there is very little causal evidence describing how the brain uses changes in neuronal spiking to render information to downstream areas (but see^59–62)^. Further work will be required to establish how changes in spiking in neuronal populations contribute to different brain computations. Nevertheless, the data presented here highlight a strong asymmetry in how rapid and sustained increases and decreases in neuronal spiking in V1 relate to perceptual reports.

### Author Contributions

JJC, NYM, and JHRM designed research. JJC, MLB, NYM, and EAP conducted research. JJC and NYM analyzed data. JJC wrote the manuscript with input from all other authors.

## Acknowledgements

The authors thank Dr. Vytas Bindokas and the University of Chicago Integrated Light Microscopy Core Facility for assistance with confocal imaging and Dr. Supriya Ghosh for assistance with electrophysiological recordings. The authors also thank Zaina Zayyad, Dr. Supriya Ghosh, and Dr. Matthew Kaufman for critical feedback on the manuscript. This work was supported by an Albert O. Beckman Postdoctoral Fellowship (JJC), the Vannevar Bush Faculty Fellowship (DJF), and NIH U01-NS090576 and NIH U19-NS107464.

## Online Methods

### Mouse Strains

All animal procedures were in compliance with the guidelines of the NIH and were approved by the Institutional Animal Care and Use Committee at The University of Chicago. Mouse lines were obtained from The Jackson Laboratory. Data come from parvalbumin-Cre mice (PV, 11 mice, 6 female; JAX stock #017320^21^, somatostatin-Cre mice (SST, 2 mice, both male; Jax stock #013044^22^, and Emx-1 Cre mice (Emx, 7 mice, 2 female, Jax stock #005628^23^. Experimental animals were heterozygous for Cre recombinase in the cell type of interest (outbred by crossing homozygous Cre-expressing strains with wild type BALB/c mice, Jax stock #000651). Mice were singly housed on a reverse light/dark cycle with ad libitum access to food. Mice were water scheduled throughout behavioral experiments, except for periods around surgeries. Mice used for electrophysiological recordings had ad libitum access to food and water.

### Cranial window implant

Mice (3–5 months old) were implanted with a headpost and cranial window to give stable optical access for photostimulation during behavior^15, 20, 50^. Animals were anesthetized with ketamine (40 mg/kg, i.p.), xylazine (2 mg/kg, i.p.) and isoflurane (1.2–2% in 100% O_2_). Using aseptic technique, a headpost was secured to the skull using acrylic (C&B Metabond, Parkell) and a 3 mm craniotomy was made over the left cerebral hemisphere (3.0 mm lateral and 0.5 mm anterior to lambda) to implant a glass window (0.8 mm thickness; Tower Optical).

### Intrinsic autofluorescence imaging

We located V1 by measuring changes in the intrinsic autofluorescence signal using visual stimuli and epifluorescence imaging^63^. Autofluorescence produced by blue excitation (470 ± 40 nm, Chroma) was collected using a green long-pass filter (500 nm cutoff) and a 1.0x air objective (Zeiss; StereoDiscovery V8 microscope; ∼0.11 NA). Fluorescence was captured with a CCD camera (AxioCam MRm, Zeiss; 460×344 pixels; 4×3 mm field of view). The visual stimuli were full contrast drifting Gabors (10° SD; 30°/s; 0.1 cycles/deg) presented for 10 s followed by 6 s of mean luminance. The response to the visual stimulus was computed as the fractional change in fluorescence during the first 8 s of the stimulus presentation compared to the average of the last 4 s of the preceding blank.

### Viral injections and ChR2 stimulation

Virus injections were targeted to a monocular region of V1 based on each animal’s retinotopic map (+25° in azimuth; between −15° to +15° in elevation). Before virus injection, mice were anesthetized (isoflurane, 1–1.5%), and the glass window was removed using aseptic technique. We used a volume injection system (World Precision Instruments) to inject 200-400 nl of AAV9-Flex-ChR2-tdTomato (∼10^11^ viral particles; Penn Vector Core) 300 µm below the pial surface. The virus was injected at a rate of 40 nl/min through a glass capillary attached to a 10 μL syringe (Hamilton). Following the injection, a new cranial window was sealed in place. Several weeks after injection, we localized the area of ChR2 expression using tdTomato fluorescence, and attached an optical fiber (400 µm diameter; 0.48 nA; Doric Lenses) within 500 µm of the cranial window (∼1.3 mm above the cortex). We delivered light though the fiber from a 455 nm LED (ThorLabs) and calibrated the total power at the entrance to the cannula. Optogenetic stimulation began no earlier than 4 weeks after injection. We prevented optogenetic stimuli from cueing the animal to respond by wrapping the fiber implant in blackout fabric (Thor Labs) that attached to the headpost using a custom mount.

### Behavioral tasks

Mice were trained to respond to changes in a visual display for a water reward using a lever while head fixed ^64^. In the primary experiment, a static 50% contrast Gabor stimulus was continuously on the screen, presented on a uniform background with the same average luminance. Mice initiated trials by depressing a lever. Following a random delay (400-3000 ms), the contrast of the Gabor stimulus either incremented or decremented (interleaved). The Gabor stimulus (SD 5-7°, 0.1 cycles/deg, odd-symmetric) changed contrast for the duration of a brief response window. The size of the contrast change varied randomly from trial to trial across a range that spanned behavioral thresholds for both increments and decrements. The mouse had to release the lever within the response window running from 100 ms to 700 or 900 ms after change onset to receive a reward. Following completion of the trial, the contrast of the Gabor stimulus returned to 50%. Stimuli for each animal were positioned at a location that corresponded to the V1 representation expressing ChR2. Early releases and misses resulted in a brief timeout before the start of the next trial. Behavioral control, data collection and analysis were done using custom software written using Objective-C, MWorks (mworks-project.org), Matlab (MathWorks) and Python.

Optogenetic stimulation did not begin until animals worked reliably for hundreds of trials each day and performance was stable at threshold for both increments and decrements. This typically required ∼2.5 months of training. During optogenetic experiments, we activated ChR2 expressing neurons on a randomly selected half of trials for a single contrast change (around ±15% for all mice). These change magnitudes were chosen for stimulation as they approximated the increment and decrement detection thresholds and thus maximized our ability to resolve an impairment or facilitation of detection capability. We aligned the opsin illumination with visually evoked spiking in V1 by delaying the optogenetic stimulus by 35 ms relative to the appearance of the visual stimulus on the monitor. Opsin illumination persisted until the end of the trial to prevent mice from using the offset of optogenetic input as a task-relevant signal. The optogenetic stimulation intensity was fixed within a session and chosen for each mouse basis based on 1 or 2 preliminary testing sessions that were not included in the main analysis. Using these preliminary observations, powers were selected to avoid saturating behavioral performance (ranges for high power sessions: Emx: 0.12-0.25 mW; PV: 0.12-0.25 mW). Following data collection at high powers, we conducted additional sessions in some mice at lower optogenetic stimulus intensities to determine how changes in performance scaled with power (ranges 0.02-0.15 mW).

Follow up experiments included retraining some PV mice (n=2; both female) to detect contrast increments of a counterphasing Gabor (2 or 4 Hz). The average contrast of Gabor stimulus was held at 20%, except during contrast changes. Changes in contrast were synchronized with zero crossings of the temporal modulation to avoid generating instantaneous luminance steps that could cue the animal to respond. Optogenetic stimulation was delivered on a random subset of trials for a moderate contrast change magnitude (+30%) and the powers used with counterphase-modulated stimuli were identical to those used in the main experiments. As above, optogenetic stimulation was delivered from stimulus onset until the end of the trial. Other task variations included shortening the duration of visual and optogenetic stimulation or ramping and stepping optogenetic stimuli up or down during contrast changes.

### Histology

Mice were perfused with 10% pH-neutral buffered formalin (Millipore Sigma Inc.), after which the brain was removed and submerged in fixative for 24 hr. The brain was subsequently rinsed with PBS, placed in a 30% sucrose PBS solution until it sank. Brains were sectioned at 40 µm on a freezing microtome, mounted and cover slipped with DAPI Fluoromount-G (Southern Biotech). tdTomato expression and DAPI labeling were visualized with 561 and 405 nm excitation light respectively, using a Leica SP5 Confocal Microscope.

### Behavioral data analysis

The proportion correct for each contrast change level was determined using trials in which the subject either responded correctly (hit) or failed to respond (miss). Trials in which the animal released the lever before stimulus onset (false alarm) were not considered in performance analyses. Sessions in which the false alarm rate or the miss rate was greater than 50% were excluded from analysis. We estimated the likelihood of observing a false hit by calculating the conditional probability that the animal would release the lever in each 100 ms bin given that a stimulus had yet to occur in that or earlier bins. The false hit rate represents the probability that the mouse would get a trial correct due to spontaneous lever releases independent of detecting stimulus changes and is thus a lower bound on performance. The false hit rate was low (Emx median 5.3%; range, 3.8-6.3%; PV median 5.3%; range, 2.8-8.3%), demonstrating that mice were relying on stimulus changes to guide their responses. To correct for false hits, we subtracted a randomly selected fraction of correct trials from each contrast level based on the estimated false hit probability observed in the corresponding session.

When performance data were fit to psychometric functions, we first corrected for the estimated false hit probability as described above. This correction was typically small (median hits removed 6.8%; range, 4.5–12.9% for 38 sessions from 4 mice). Corrected performance data were then fit with a Weibull cumulative distribution function using non-linear least squares and variance weighting of each mean. The two psychometric functions (with and without ChR2 stimulation) were fit simultaneously using four parameters: individual thresholds (α_unstimulated_, α_stimulated_), a common lapse rate (γ), and a common slope (β) such that:

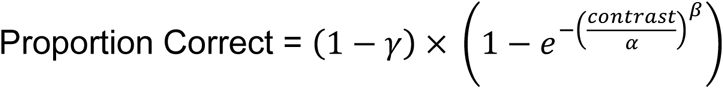

Threshold confidence intervals were estimated using a bootstrap (1000 repetitions, p < 0.05, one-tailed).

To compare how optogenetic increments and decrements affected lever responses relative to the trial-time-matched false rate (Figure 3), we used the stimulus onset times for 0% contrast change trials in which mice correctly responded to optogenetic stimuli. We restricted our analyses to stimulus onset times that occurred before the final 400 ms of all possible onset times as the small number of observations for the longest trial times made this calculation unstable over the response window. Using only trials in which stimuli had yet to occur by the stimulus onset time, we expressed the lever release time for false alarms relative to the stimulus onset time. Thus, the false alarm distribution aligned to the onset of stimuli served as a time-in-trial matched measure for the lever release probability over time for static optogenetic input. This process was repeated within each session and the probability of lever releases were compared for optogenetic increments, decrements, and time-matched false alarms.

### Electrophysiological recordings

We recorded extracellularly from V1 in awake, head-fixed mice (n=4 Emx, 1 female; n=4 PV, 3 female) using multisite silicon probes (Neuronexus, Inc.; 32-site model 4 × 8-100–200-177). Some mice were first used for behavioral experiments (2 PV, both female), while the rest were untrained but injected with opsins before recording. Electrodes were electroplated with a gold solution mixed with carbon nanotubes^65, 66^ to impedances between 200-500 kΩ.

At the start of recording sessions, mice were anesthetized with isoflurane (1.2–2% in 100% O_2_), placed in a sled and head-fixed. While anesthetized, the eyes were kept moist with 0.9% saline. We visualized ChR2-expressing areas of monocular visual cortex by imaging tdTomato fluorescence with a fluorescence microscope and camera (Zeiss, Inc). The cranial window was then removed and the electrodes lowered through a slit in the dura. We then positioned an optic fiber above the cortex at a distance comparable to that used during behavioral experiments (1.0-1.5 mm). The craniotomy was then covered with 3% agarose dissolved in aCSF (Millipore Sigma Inc., TOCRIS respectively) Following the recovery period of 1 hour, anesthetic was removed and we waited at least an additional hour for recovery from anesthesia before recording.

The electrode was advanced to locate responsive units, and was allowed to settle for 30 minutes before collecting data. Delivery of visual and optogenetic stimuli and data acquisition was computer controlled. Concurrent visual and optogenetic stimuli matched those used during behavioral experiments except that visual and optogenetic changes were presented for 500 ms rather than 700-900 ms and the visual stimuli filled the video display. We recorded at least 25 repetitions of each stimulus condition in a given stimulus set. For optogenetic stimuli presented in isolation (Figure 3), optogenetic input ramped up for 250 ms at the beginning of each stimulus to a moderate baseline power (0.5 mW), where it remained for 750 ms, and stepped up or down in intensity (randomly interleaved) for 250 ms before returning to the baseline for the remainder of the trial.

Electrode signals were amplified, bandpass filtered (750 Hz to 7.5 kHz) sampled around threshold crossings (Blackrock, Inc.) and spikes were sorted offline (OfflineSorter, Plexon, Inc.). Visually responsive units were taken as those with a 10% change in the average firing rate during the 50-250 ms after stimulus onset (stimulus period) relative to the average firing rate during the baseline epoch (baseline period −250 to −50 ms before stimulus onset) for the largest stimulus intensities. Optogenetically responsive units were defined as any unit with a significant difference (p < 0.05; signed-rank test) in firing rate during the stimulus period for the same visual stimulus with and without optogenetic stimulation. For optogenetic stimuli presented in isolation (Figure 6), optogenetically responsive units were classified based on significant changes in firing for optogenetic increments or decrements (from 0-250 ms following optogenetic steps) relative to pre-change firing rates (from −250 - 0 ms; *P* < 0.05; signed-rank test). We recorded both single and multi-units but did not differentiate between them because our primary interest was how optogenetic manipulations affect visually evoked responses across the V1 population.

### Neural network models

We trained recurrent neural network (RNN) models on a task similar to the one that the mice perform to test whether *in silico* networks adopt the same strategies as *in vivo* networks. RNNs were trained and simulated using the Python machine learning framework TensorFlow^67^, and the network architecture was based on our previous study^68^. Briefly, all networks consisted of orientation and contrast selective input neurons (whose firing rates are represented as ***u***(*t*)) that projected onto 100 recurrently connected neurons (whose firing rates are represented as ***h***(*t*), which in turn projected onto the output layer (Figure 5). Recurrently connected neurons never sent projections onto themselves.

The activity of the recurrent neurons was modeled to follow the dynamical system^69^:

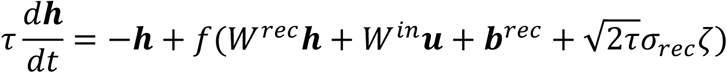

where τ is the neuron’s time constant (set to 50 ms), *f*(·) is the activation function, *W^rec^* and *W^in^* are the synaptic weights between recurrent neurons, and between input and recurrent neurons, respectively, ***b**^rec^*is a bias term, ζ is independent Gaussian white noise with zero mean and unit variance applied to all recurrent neurons, and *σ_rec_* is the strength of the noise (set to 0.05). To ensure that neuron’s firing rates were non-negative and non-saturating, we chose the rectified linear (ReLu) function as our activation function: *f*(*x*) = *max*(0, *x*).

To simulate the network, we used a first-order Euler approximation with time step Δt:

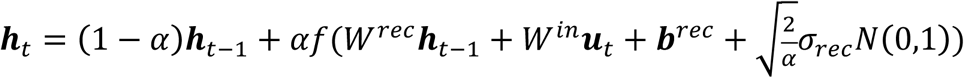

where 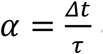 and *N*(0,1) indicates the standard normal distribution.

The decision to release the lever (when there was a contrast change) or to hold the lever (when there was no contrast change) was mediated by a competition between two output units. The 80 excitatory neurons linearly projected onto the output unit associated with releasing the lever:

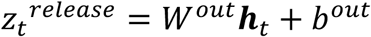

where *W^out^* are the synaptic weights between the excitatory neurons and the output unit, and *b^out^* is a bias term. The activity of the output unit associated with holding the lever was simply the negative of the activity of the unit associated with releasing the lever: 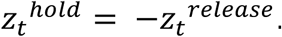

We then calculated the network policy, π*_t_*, which was the probability of holding or releasing the lever, by taking the softmax of these two values:

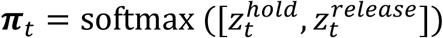

To maintain separate populations of 80 excitatory and 20 inhibitory neurons, we decomposed the recurrent weight matrix, *W^rec^* as the product between a matrix for which all entries are non-negative, *W^rec^*^+^ whose values were trained, and a fixed diagonal matrix, *D*, composed of 1s and −1s, corresponding to excitatory and inhibitory neurons, respectively^69^:

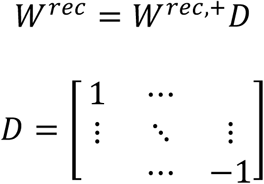

Initial connection weights between excitatory neurons were randomly sampled from a Gamma distribution with shape parameter of 0.1 and scale parameter of 1.0, and then multiplied by 0.25. Initial connections weights projecting to or from inhibitory neurons were sampled from a Gamma distribution with shape parameter of 0.1 and scale parameter of 1.0 and then multiplied by 0.5. Initial bias values were set to 0.

Networks consisted of 24 orientation and contrast selective input neurons. The tuning of the input neurons followed a Von Mises’ distribution, such that the activity of the input neuron was

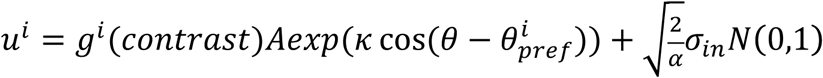

where θ is the orientation of the stimulus (always fixed at 0°), 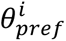 is the preferred direction of input neuron *i*, κ was set to 2, and *A* was set to 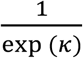. The strength of the input activity noise, *σ_in_*, was set to 0.05. The function *g^i^*(*contrast*) determined how neuron *i* responded to different contrasts. Half (12) of the input neurons were contrast increasing, defined as *g^i^*(*contrast*) = *contrast*, while the other half were contrast decreasing, defined as as *g^i^*(*contrast*) = 1/*contrast*.

### Contrast change detection task for network model

The networks were trained to indicate whether the stimulus contrast changed by responding within a fixed interval. Trials lasted 3000 ms, divided into 10 ms steps. A Gabor patch with an orientation of 0° and a baseline contrast level was presented from the start of the trial, and at a random time, the contrast either doubled or was halved for 100 ms, before returning to baseline. The time of the contrast change was randomly sampled from an exponential distribution with a time constant of 1300 ms, plus 400 ms. If the contrast change did not occur before the end of the trial, the network was rewarded for maintaining hold of the bar throughout the trial. The network received a reward of +1 if it chose to release the lever during the 100 ms duration contrast change, a reward of – 0.1 if it chose to either release the lever before the contrast change (i.e. false alarm), or not release during the contrast change (i.e. miss).

### Network training

The RNNs using the actor-critic reinforcement learning method^70^, in which the networks were trained to maximize the discounted cumulative future reward:

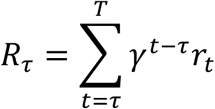

where *γ* ∈ (0,1] is the discount factor and *r_i_* is the reward given at time *t*. The network was trained to estimate this discounted future reward as a linear projection from the recurrent units

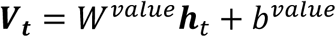

by minimizing the loss function:

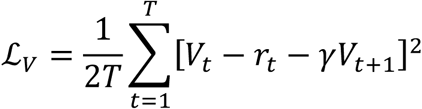

Concurrently, the network adjusts the network policy, π*_i_* (described above) to select the actions that would lead to the greatest cumulative reward by minimizing the loss function:

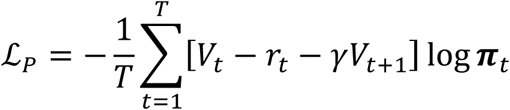

The network was also encouraged to explore different strategies by maximizing the entropy of the policy output:

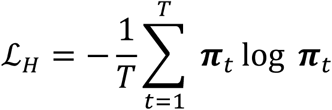

Finally, the network was encouraged to solve the task using low levels of neural activity by minimizing the L2 norm of the recurrent neuron firing rates:

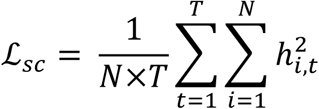

where *h_i,t_* is the neural activity of the ith recurrent neuron at time t.

All together, the overall loss function is the weighted sum of all four terms:

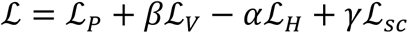

where *α* and *β* were set to 0.01. To understand different network solutions across various metabolic constraints, *γ* was randomly sampled for each network from a logarithmically uniform distribution between 10^-6.5^ and 10^-0.5^.

We trained all network parameters using the Adam version of stochastic gradient descent, with 1st and 2nd moment decay rates set to their default values (0.9 and 0.999, respectively). All networks were trained for 50,000 batches, with a batch size of 1024 trials and a learning rate of 0.001.

### Analysis of Neural Network Responses

To link unit responses with network decisions to respond or withhold responses to contrast changes, we calculated the mean normalized ROC value for each network. First, we calculated the ROC for each excitatory unit by comparing its firing rate distributions when the network decided to release versus continue holding the lever following contrast changes. The ROC values for increments and decrements were then averaged for each unit.

Next, we summed a rescaled version of each units ROC value, normalized by the absolute value of this metric, such that:

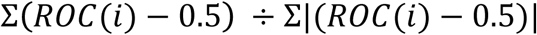

Where *i* is the index of all units in the network. Here, a value of −1 is the extreme case where all units in the network have lower firing rates when the network releases than when it withholds, whereas a value of +1 indicates all units had higher firing rates when the network responds compared to trials in which it does not respond.

## Data Availability

Data that support the findings of this study are available from the corresponding author upon reasonable request.

## Code Availability

Custom code required for experimental control is available from the corresponding author upon reasonable request.

