## Supplemental Data for "Perceptual detection depends on spike count integration"

**Cone et al. Supplemental Information**

**Table S1**

| **Emx-Cre** | **Decrements** | | | **Increments** | | |
|---|---|---|---|---|---|---|
| **Mouse** | **Hits/Miss unstim** | **Hits/Miss stim** | **p-value** | **Hits/Miss unstim** | **Hits/Miss stim** | **p-value** |
| **1** | 79/73 | 153/31 | $< 10^{-9}$ | 69/78 | 154/32 | $< 10^{-11}$ |
| **2** | 82/121 | 166/123 | $< 10^{-3}$ | 74/110 | 168/199 | $< 10^{-3}$ |
| **3** | 66/86 | 149/87 | $< 10^{-3}$ | 56/99 | 123/74 | $< 10^{-6}$ |

**Table S1. Summary of changes in detection performance produced by optogenetic stimulation of pyramidal neurons in Emx-Cre mice.** We combined data across sessions within individual mice and calculated the relative proportion of hits and misses separately for contrast changes (±15%) with and without optogenetic stimulation of pyramidal neurons. Cumulative data from the three Emx mice is reported above, numbers correspond to the total counts of hits/misses for contrast decrements (left) and increments (right). P-values correspond to the results of Fisher's exact test comparing the proportions of hits/misses with and without stimulation independently for contrast decrements and increments. Optogenetic stimulation of excitatory neurons similarly facilitated contrast change detection, regardless of sign, in all three tested mice.

**Table S2**

| PV-Cre | Decrements | | | Increments | | |
|---|---|---|---|---|---|---|
| **Mouse** | **Hits/Miss unstim** | **Hits/Miss stim** | **p-value** | **Hits/Miss unstim** | **Hits/Miss stim** | **p-value** |
| **1** | 295/113 | 77/132 | $< 10^{-23}$ | 203/83 | 92/82 | $< 10^{-16}$ |
| **2** | 181/222 | 14/199 | $< 10^{-25}$ | 149/255 | 19/195 | $< 10^{-15}$ |
| **3** | 76/95 | 31/95 | $< 10^{-7}$ | 69/47 | 30/88 | $< 10^{-6}$ |
| **4** | 82/81 | 28/133 | $< 10^{-9}$ | 74/90 | 34/129 | $< 10^{-5}$ |
| **5** | 77/99 | 41/118 | $< 10^{-4}$ | 74/86 | 43/116 | $< 10^{-3}$ |
| **6** | 178/94 | 62/184 | $< 10^{-19}$ | 197/79 | 97/149 | $< 10^{-12}$ |

**Table S2. Summary of changes in detection performance produced by optogenetic stimulation of PV interneurons in PV-Cre mice.** As was done for Emx mice, we combined data across all sessions at the tested contrast changes (±15%) and calculated the relative proportion of hits and misses separately for trials with and without optogenetic stimulation of PV interneurons. Cumulative data from the six PV mice is shown above, numbers correspond to the total counts of hits/misses for contrast decrements (left) and increments (right). P-values are the results of Fisher's exact test comparing the proportions of hits/misses with and without stimulation independently for contrast decrements and increments. Optogenetic stimulation of PV interneurons similarly impaired detection of decrements and increments in all six tested mice.

### Figure S1

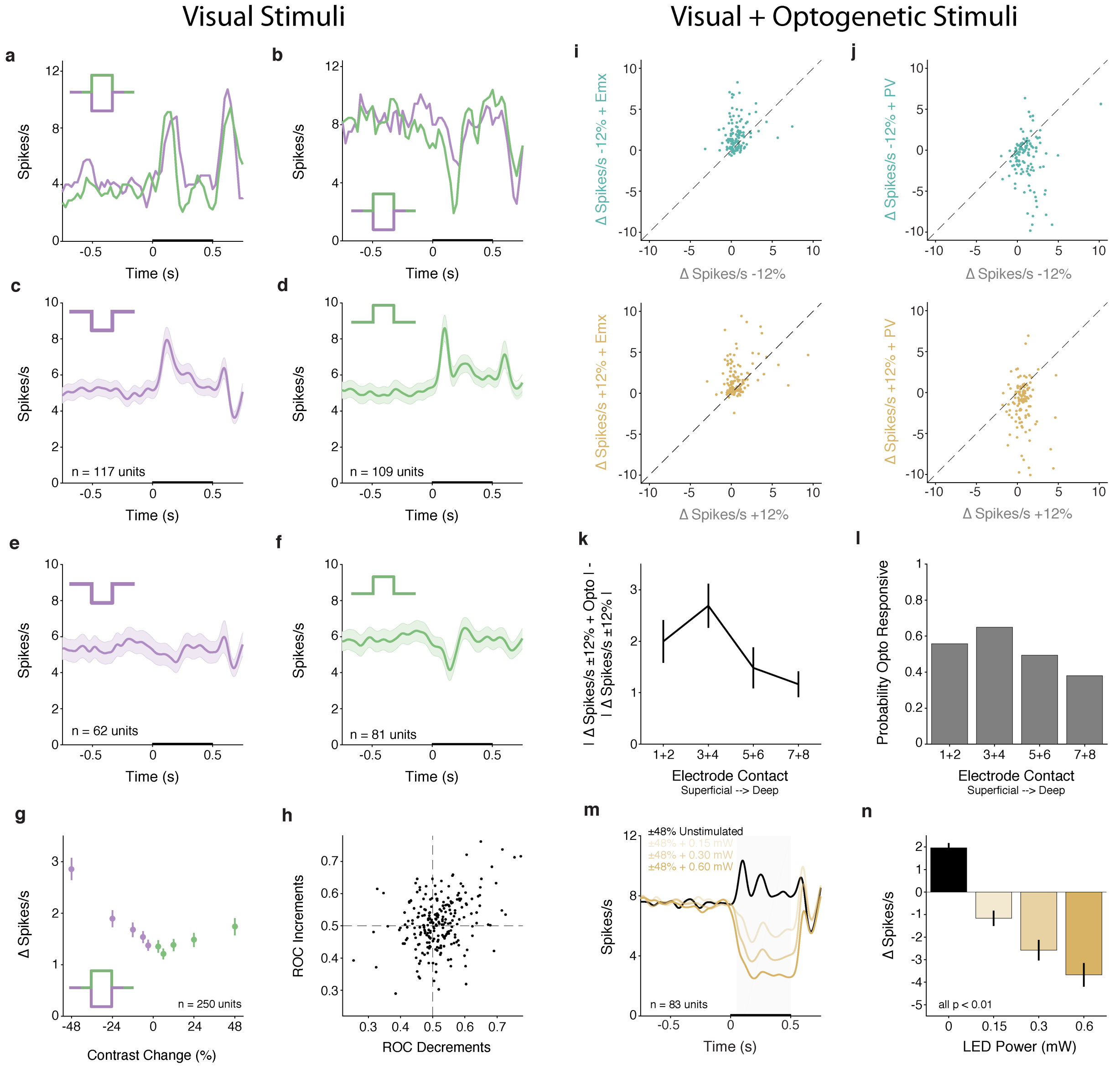


**Figure S1. The V1 population exhibits diverse responses to increments and decrements in contrast, while the effects of optogenetic excitation and inhibition are consistent with expectations and scale by both depth and power.** A) Average contrast change responses from a representative unit that was excited by halving (decrements, purple) and doubling (increments; green) the stimulus contrast. Legend depicts the contrast profile. Visual stimulus duration is indicated by the thickening of the x-axis in all PSTHs. Bin size = 25 ms, smoothed. B) Same as in A except for a unit that was inhibited by contrast changes.

C-G) Our primary goal was to assess the total stimulus-evoked signal present in V1, we classified units as excited or inhibited if the average firing rate increased or decreased by 10% relative to baseline when the initial change in contrast either halved or doubled. Using this criterion, many units were excited by contrast decrements (C; 117/250, 47%) and many by contrast increments (D; 109/250, 44%). Smaller proportions of units were inhibited by decrements (E; 62/250, 25%) or by increments (F; 81/250, 32%). In addition to being less prevalent, inhibitory responses were also weaker compared to excitatory responses, especially for contrast decrements. C) Gaussian filtered ($\sigma$ = 25 ms) PSTH for units that were excited by contrast decrements (n = 117 units). Shaded region = SEM. D) Same as in C except for units excited by contrast increments (n = 109 units). E-F) Same as in C, D except for units that were inhibited by contrast decrements (E, n = 62 units) or contrast increments (F, n = 81 units). G) Evoked change in firing rate relative to baseline across the population for all contrast changes. H) To the measure the ability of individual units to detect visual contrast changes, we performed an ROC analysis for each unit that compared the pre-stimulus (-250 - -50 ms before stimulus onset) with stimulus-evoked firing rates (50-250 ms after stimulus onset) evoked by a halving (decrements, x-axis) or doubling (increments, y-axis) of the baseline contrast. Points depict the ROC value for discriminating between the average baseline and stimulus firing rates for decrements (x-axis) and increments (y-axis) in contrast. While the median ROC values were both near 0.5 (decrements = 0.52; increments = 0.51), the range of ROCs values suggested there was reasonable predictive power for firing rate decreases (ROC<0.5) or increases (ROC>0.5) for either contrast change type (decrements: range 0.29-0.85; increments: range 0.25-0.79).

I) Optogenetic stimulation (0.3 mW) potentiates contrast change responses in Emx-Cre mice. Scatter plot shows the evoked change in firing rate (stimulus epoch – baseline) for each unit recorded in Emx-Cre mice (n = 119 units) in response to moderate contrast decrements (top) without (x-axis) and with (y-axis) optogenetic stimulation of pyramidal neurons. 83 of 119 (70%) units had higher evoked responses for trials with optogenetic stimulation. Moreover, the distributions of firing rate changes were significantly different (decrements median = +0.56 ΔSpikes/s, 0.08-1.09 IQR; visual + optogenetic trials median = +1.4 ΔSpikes/s, 0.37-2.76 IQR, $p < 10^{-7}$, Komolgorov-Smirnov test). Similar effects were observed for contrast increments (bottom) as 81/119 (68%) of units had higher evoked responses for contrast increments when they were paired with optogenetic stimulation. The distributions of firing rate changes were significantly different from one another (increments median = +0.39 ΔSpikes/s, -0.07-1.07 IQR; visual + optogenetic trials median = +1.23 ΔSpikes/s, 0.33-2.6 IQR, $p < 10^{-4}$, Komolgorov-Smirnov test). J) Same as in I except for PV mice (n = 131 units). 105 of 131 (80%) units had lower evoked responses for trials with optogenetic stimulation and the distributions of firing rate changes were significantly different (decrements median = +1.04 ΔSpikes/s, 0.25-1.83 IQR; visual + optogenetic trials median = -0.66 ΔSpikes/s, -2.9-0.48 IQR, $p < 10^{-13}$, Komolgorov-Smirnov test). Similar effects were observed for contrast increments (bottom). 102 of 131 (78%) of units had lower evoked responses for contrast increments paired with optogenetic activation of PV interneurons compared to the same increment without optogenetic stimulation. (increments median = +0.66 ΔSpikes/s, 0.19-1.13 IQR; visual + optogenetic trials median = -0.94 ΔSpikes/s, -3.45-0.29 IQR, $p < 10^{-16}$, Komolgorov-Smirnov test).

K-L) The strength of optogenetic modulation differs by recording depth. We examined how the effects of optogenetic stimulation on population responses varied across recording sites located at different depths. For our recordings, we used 4 shank probes with 8 contacts per shank (A32 4x8, NeuroNexus Inc.) and recording sites were separated by 100 μm. A straightforward proxy for depth is to separate data from superficial versus deep electrode contacts. We calculated the absolute change in spike rate during the stimulus window (+50 - 500 ms) relative to the pre-stimulus period (-500 to -50 ms) for ±12% contrast changes (increments and decrements). This was done separately for trials with and without optogenetic stimulation. Using the absolute change in spike rate allowed us to combine data from PV-Cre and Emx-Cre mice as the effects of optogenetic stimulation on neuronal activity were opposite in sign. We then subtracted the absolute change in spiking on trials without optogenetic stimulation from the response on trials with stimulation to isolate the contribution of optogenetic input to spiking during the stimulus period. Thus, each unit contributed two data points in this dataset (contrast increments, decrements). This metric captures the absolute difference in spike rate induced by optogenetic stimulation above what would be expected based on the visual stimulus alone for each unit (n = 250 units, 4 Emx-Cre, 4 PV mice). The data were then grouped based on which contact the units were recorded on, starting with the most superficial (1+2) to the deepest (7+8). As would be expected from light scattering through tissue, optogenetic effects on unit responses were strongest for the most superficial contacts and appeared to fall off with increasing depth (mean ΔSpikes/s: sites 1&2: 2.0, 0.42 SEM; sites 3&4: 2.7, 0.43 SEM; sites 5&6: 1.48, 0.28 SEM; sites 7&8: 1.17, 0.25 SEM). K) Plot shows the mean±SEM difference in evoked response with versus without optogenetic input for all units recorded at different depths. We found that the strength of optogenetic modulation significantly depended on depth ($p < 0.001$, Kruskal-Wallis test). *Post hoc* comparisons showed the optogenetic modulation was significantly greater for sites 3&4 (versus 5&6 and 7&8, both $p < 0.01$; trend for site 1&2, $p = 0.07$; Dunn–Šidák correction). L) The depth related differences in the strength of optogenetic modulation was in close agreement with the probability of recording units significantly modulated by the LED. Units were classified as optogenetically responsive if their firing rate during the stimulus period on trials with optogenetic stimulation was significantly different from its' firing rate during the stimulus window on trials without optogenetic input ($p < 0.05$, Wilcoxon signed rank test, see Main Text). The probability of recording from an optogenetically responsive unit differed by depth (sites 1&2: 56%, 29 of 52 units; sites 3&4: 65%, 24 of 37 units; sites 5&6: 50%, 38 of 77 units; sites 7&8: 38%, 32 of 84 units). This pattern of results is consistent with expectations. In addition to the contribution of light scattering with depth, our viral injections were targeted to between 250-400 μm from the cortical surface. Moreover, PV interneurons are most concentrated in layer 4[1] while Emx-Cre mice generate widespread transgene expression in all layers below layer 1[2]. Together, these observations suggest our optogenetic manipulations most strongly affected responses in superficial cortical layers (likely II/III) but units were affected by optogenetic stimulation throughout cortex.

M-N) In a subset of electrophysiology sessions in PV mice (n=3; 12 recording sites), we collected additional datasets to assess whether visually evoked changes in firing rate across the population depended on power used for PV interneuron activation. In these datasets, we presented only the largest contrast changes (±48% contrast changes from a 50% contrast static Gabor) so as to elicit the strongest visual responses. In addition to trials without optogenetic stimulation, we also presented visual stimuli concurrently during optogenetic activation of PV neurons. We tested 3 different stimulation powers (0.15, 0.3, 0.6 mW), the highest of which was ~2x greater than the highest power used in behavioral experiments shown in Figure 2. Visual and optogenetic stimuli were presented for 500 ms. M) Population PSTH (n=83 units) combined for both contrast change types (±48%, each neuron contributes both an increment and decrement response to each trace while the color scheme follows the increment convention used in the manuscript). Spikes were convolved with a Gaussian with $\sigma$ = 25 ms. Black line depicts the population response without optogenetic activation of PV interneurons. Darkness of the colored lines depicts the population response for different PV stimulation powers. N) PV-mediated suppression of firing scales with power. Bar plot compares the change in firing rate from +50 to +500 ms after stimulus onset relative to the pre-stimulus period (-500 to -50 ms). The strength of PV-mediated suppression significantly depends on LED power ($p < 10^{-34}$, Friedman's test; post hoc all comparisons $p < 0.01$, Dunn–Šidák correction). These observations held when increments or decrements were tested in isolation (Decrements: $p < 10^{-15}$, post hoc: 0.15 mW versus 0.3 mW $p > 0.05$, otherwise all other comparisons $p < 0.01$; Increments: $p < 10^{-16}$, *post hoc*: 0.3 mW versus 0.6 mW $p > 0.05$, otherwise all other comparisons $p < 0.05$; both Friedman's test with Dunn–Šidák correction).

**Figure S2**

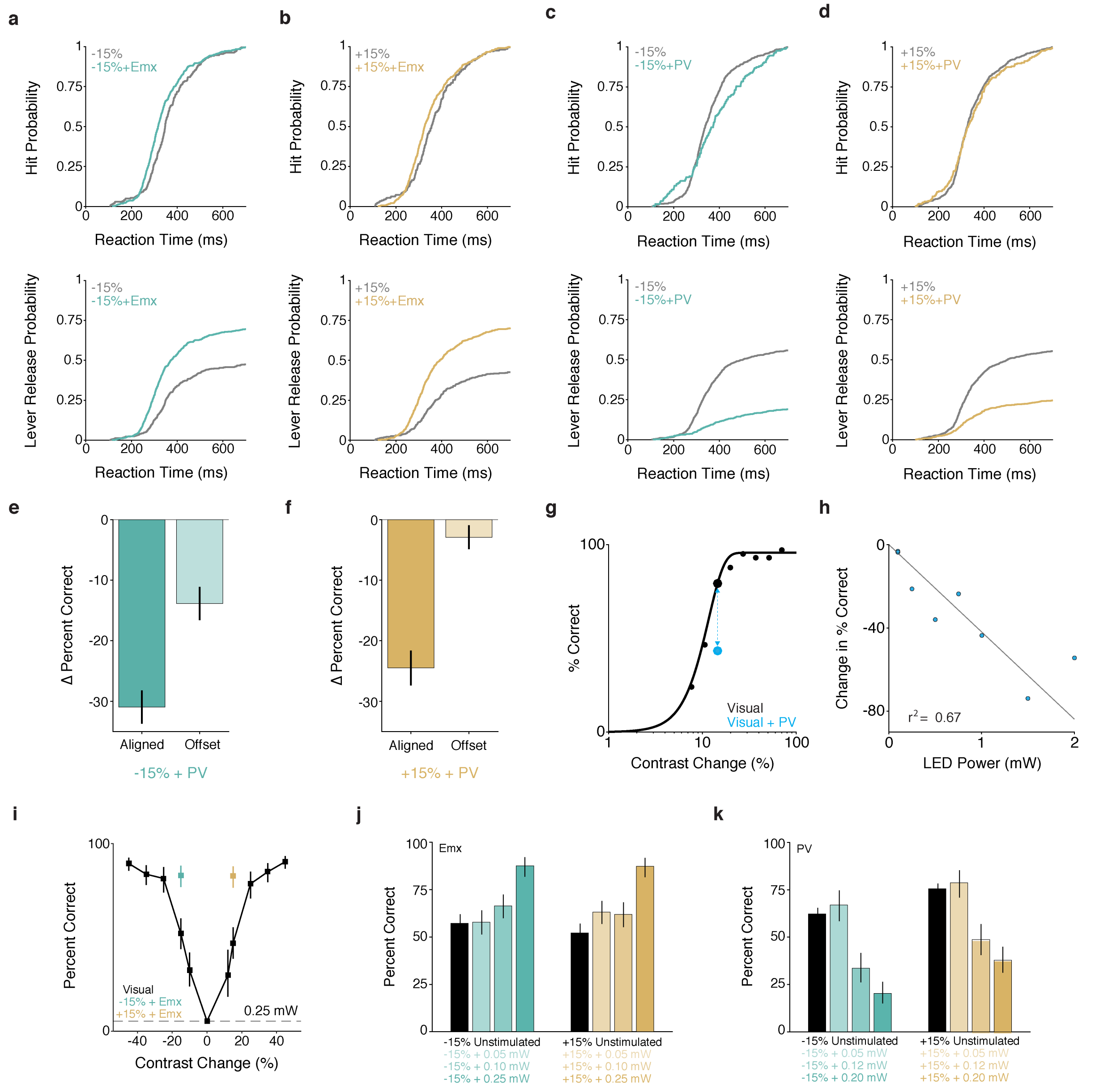


**Figure S2. The effect of optogenetic stimulation on behavior depends on retinotopic alignment with the visual stimulus and stimulation power.** The processing time required to detect and respond to a visual stimulus strongly depends on the stimulus intensity[3,4]. We thus sought to examine effects of optogenetic stimulation on reaction times. We combined reaction time data across all mice (n=3) at the tested contrast changes to determine if the time of lever releases following stimulus onset was affected by optogenetic stimulation. First, we only examined trials in which the mice correctly detected stimuli (A-B, top, hits). We found that optogenetic stimulation of excitatory neurons in Emx-Cre mice significantly enhanced reaction times for both contrast decrements (A, top; unstimulated (gray line) median = 347 ms, 296-414 IQR; stimulated (aqua line) median = 316 ms, 273 – 388 IQR; $p < 10^{-4}$;) and increments (B, top; unstimulated (gray) median = 356 ms, 295 – 424 IQR; stimulated (gold) median = 326 ms, 276 – 409 IQR; $p < 0.01$; Komolgorov-Smirnov test) on hit trials. Note: the absence of responses before 100 ms is due to the fact we only counted responses as hits if the lever release occurred 100 ms after stimulus onset, as otherwise responses would have been too early to be driven by visual stimuli. We also plotted reaction time data for hits and misses combined (A-B, bottom plots). These plots capture the combined effects of optogenetic stimulation on response probability (e.g., the difference in the cumulative probability of responses at the end of the reaction time window) along with the enhancement in reaction time for hit trials (the leftward shift and change in the rising phase of the lever response probability). C-D) Same as in A,B except for PV-Cre mice. PV stimulation augmented the reaction times on hit trials for contrast decrements (C, top: unstimulated (grey line) median = 338 ms, 290 – 406 IQR; stimulated (aqua line) median = 373 ms, 293 – 474 IQR; $p < 0.01$; Komolgorov-Smirnov test). However, we did not find a difference between the stimulated and unstimulated reaction time distributions for successfully detected contrast increments (D, top: unstimulated (gray) median = 326 ms, 284 - 396 IQR; stimulated (gold) median = 330 ms, 278 - 404 IQR; $p = 0.45$; Komolgorov-Smirnov test). We then examined the total response probabilities by including both hits and misses (bottom). PV stimulation strongly suppressed the probability of lever releases throughout the reaction time window.

E-F) The effect on PV stimulation on task performance depends on the alignment between the visual stimulus and the inhibited patch of visual cortex. In a subset of PV mice (n=3), we collected additional behavioral sessions (n=22) in which we moved the visual stimulus away from the retinotopic location of optogenetic stimulation. In these additional sessions (n=22 sessions), we offset the stimulus by 15-20° from the location used for primary data collection (n=25 sessions) while the LED power remained the same. Thus, despite comparable levels of PV activation, offsetting the visual stimulus should attenuate suppression of stimulus-evoked activity in V1. Indeed, optogenetic perturbations produced larger changes in performance when the visual stimulus was aligned with the stimulated patch of V1 (ΔPercent Correct = stimulated – unstimulated). This was true for both decrements (E, median ΔPercent Correct, Aligned: -31.2%, -0.41 - -0.17 IQR; Offset: -11%, -0.21 - -0.03 IQR, $p < 10^{-4}$; left) as well as increments (F, median ΔPercent Correct, Aligned: -22%, -0.35 - -0.13 IQR; Offset: -0.02%, -0.09 - 0.05 IQR, $p < 10^{-5}$; right). Thus, the ability of PV stimulation to reduce behavioral responses depends on alignment between the optogenetic stimulus and the visual representation in V1. This argues against the possibility that PV stimulation is somehow affecting performance by distracting the animal or impairing motor planning or execution.

(G) Representative behavioral performance from a single contrast decrement session where one brief contrast decrement (100 ms) was paired with stimulation of PV interneurons. Optogenetic power of 0.5 mW was the stimulation intensity used in this example session. Filled dots represent performance for trials with (blue) and without (black) activation of PV interneurons as the magnitude of contrast decrement is varied. Solid line depicts performance on visual only trials fit with a Weibull function. Blue dashed line connecting dots shows magnitude of perceptual impairment on the selected contrast decrement with (blue) without (black) optogenetic activation of PV neurons. (H) The perceptual impairment induced by PV activation scales with optogenetic stimulation intensity. Individual points depict performance impairment from individual sessions (note: there are two sessions at 0.1 mW). Gray line is a maximum likelihood linear regression anchored at the origin ($r^2$ = 0.67).

I-J) Data come from an Emx-Cre mouse trained to detect interleaved increases and decreases in contrast. We tested whether the effects of excitatory neuron activation on contrast perception depends on the stimulation power. I) Cumulative psychometric function depicts the probability of detection across a range of contrast changes. A random subset of ±15% contrast changes were paired with optogenetic stimulation (0.25 mW) of excitatory neurons. Different powers were used in different testing sessions (0.05 mW (8 sessions), 0.15 mW (8 sessions), 0.25 mW (7 sessions)). We combined the data across all sessions for the tested contrast changes and performed a logistic regression that compared the probability of hits as a function of stimulation power independently for increments and decrements. We found that the probability of successfully detecting either increments or decrements in contrast was significantly influenced by the power used for optogenetic stimulation (J, Decrements hit rate ± 95%

CI: 0 mW = 54%, 48-59%; 0.05 mW = 58%, 51-63%; 0.15 mW = 66%, 60-72%; 0.25 mW = 83%, 77-88%; t-statistic = 5.47, $p < 10^{-7}$; Increments hit rate ± 95%: 0 mW = 49%; 43-55%; 0.05 mW = 61%, 55-67%; 0.15 mW = 62%, 56-68%; 0.25 mW = 82%, 77-88%: t-statistic = 5.48, $p < 10^{-7}$). Thus, the degree to which optogenetic stimulation of excitatory neurons improves detection of both increments and decrements in contrast depends on the power used for stimulation.

K) Same as in J except for a PV-Cre mouse trained to detect interleaved increases and decreases in contrast. Different powers were used on different testing sessions (0.05 mW (7 sessions), 0.12 mW (7 sessions), 0.2 mW (8 sessions)). As was done in J, we performed a logistic regression that compared the probability of hits as a function of stimulation power independently for increments and decrements. We found that the probability of successfully detecting either increments or decrements in contrast was significantly influenced by the power used for optogenetic stimulation (Decrements hit rate ± 95% CI: 0 mW = 60%, 57-64%; 0.05 mW = 64%, 55-72%; 0.12 mW = 33%, 25-41%; 0.2 mW = 21%, 16-27%; t-statistic = -10.5, $p < 10^{-25}$; Increments hit rate ± 95% CI: 0 mW = 73%, 70-76%; 0.05 mW = 76%, 68-83%; 0.12 mW = 47%, 39-55%; 0.2 mW = 37%, 30-44%; t-statistic = -8.34, $p < 10^{-16}$). Thus, the magnitude of the impairment produced by PV-mediated inhibition scales with power.

### Figure S3

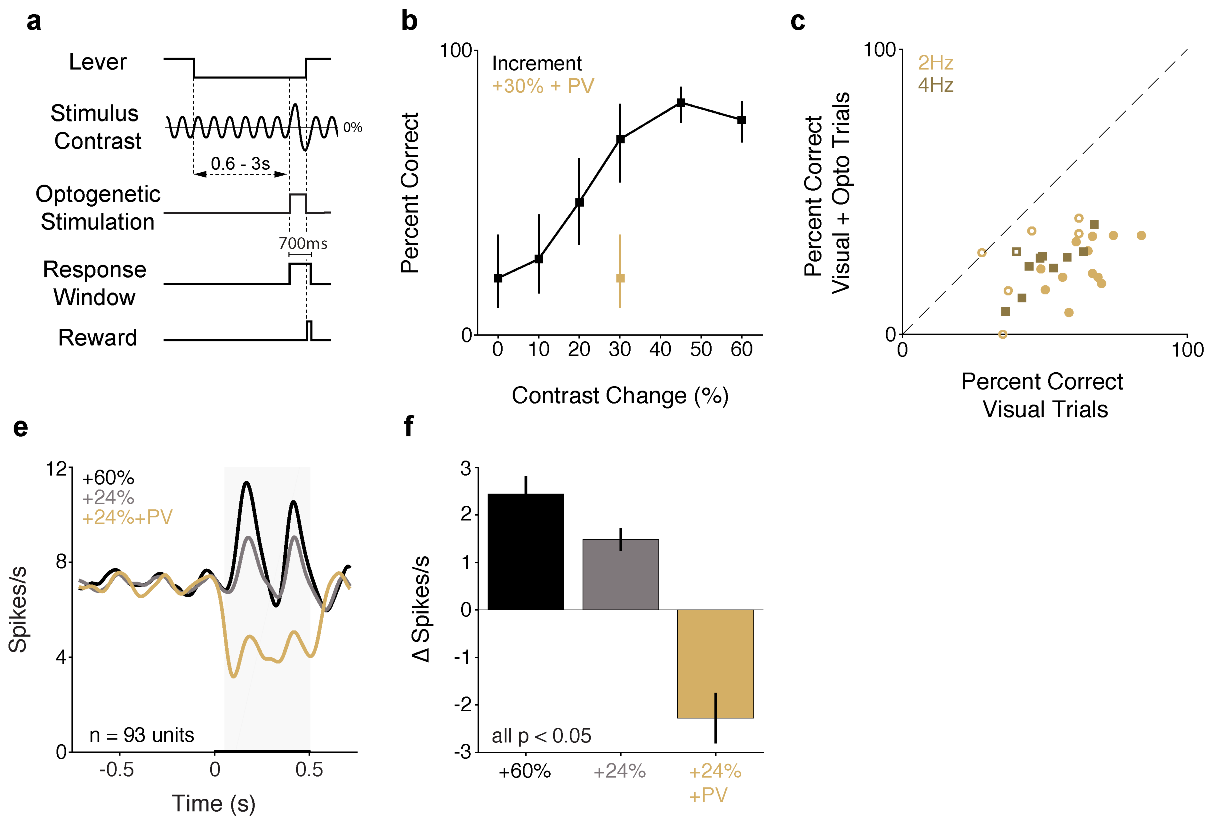


**Figure S3. Optogenetic stimulation of PV interneurons impairs detection of contrast increments presented on a counterphase modulated Gabor.** We sought to test if the effects of PV stimulation on performance depended on visual adaptation. In our main experiment, a static 50% Gabor stimulus was always present on the video display, which can attenuate visual responses due to adaptation[5–7]. We trained PV mice (n=2; both female; 1 was retrained after completing the main experiment) to detect contrast increments of a Gabor stimulus that was counterphase modulated (2 or 4 Hz).

(A) Trial schematic. The average contrast of the Gabor stimulus was held at 20%. Contrast changes were synchronized with the zero crossings of the phase modulation and were randomly selected from values spanning threshold. We reasoned that the counterphase modulation would drive the V1 population into higher rates of firing, therefore providing a larger pedestal on which to inhibit neuronal activity. We optogenetically activated PV interneurons on a random half of trials for a moderate contrast increment (+30%). As before, optogenetic stimulation was delivered starting at stimulus onset through the end of the trial. (B) Representative single session performance (hit rate ± 67% CI) from a mouse trained to detect contrast increments of counterphase modulated stimuli. Across all sessions (2 mice, 28 sessions; for 2Hz (n=18 sessions) and 4Hz (n=10 sessions), the proportion of trials in which mice successfully detected a 30% contrast increment was significantly lower on trials with optogenetic stimulation of PV neurons (median = 57.1% versus 26.9; $p < 10^{-5}$, Wilcoxon signed rank test). Performance was impaired for either counterphase modulation rate when tested in isolation (2 Hz: median = 61.5% versus 25.7, $p < 10^{-3}$; 4 Hz: median = 47.8% versus 26.7 $p < 0.01$; both Wilcoxon signed rank test). For both mice, we found a significant decrease in the cumulative proportion of hits relative to misses on trials with PV stimulation (Table S3; 2 Hz: both mice at least $p < 10^{-11}$; 4 Hz: both mice at least $p < 10^{-10}$; all Fisher's exact test) (C) Symbols depict the percent correct in individual behavioral sessions (2 mice, 28 total sessions) with (y-axis) and without (x-axis) optogenetic stimulation, separately for sessions in which the counterphase modulation frequency was 2 Hz (gold circles; 18 sessions) or 4 Hz (brown squares; 10 sessions). Filled circles indicate a significant change in detection performance (21/28 sessions; 2 Hz: 12/18 sessions $p < 0.05$; 4 Hz: 9/10 sessions $p < 0.05$; Fisher's exact test).

(D) We recorded neuronal activity in passively viewing, awake mice while presenting contrast increments on a background of a 20% contrast counterphase modulated (2 Hz) Gabor stimulus. We recorded a total of 93 units across 9 sites in 3 PV mice. Optogenetic stimulation was delivered on a near-threshold contrast change (+24%) concurrently with the onset of the visual stimulus. Modulating the stimulus generated a higher pedestal of spontaneous and stimulus-evoked activity in V1. We calculated the average firing rate during the 500 ms preceding the onset of the visual stimulus across all recorded units when mice viewed either counterphasing or static stimuli. Baseline firing rates were significantly elevated for a 20% contrast counterphase modulated stimulus compared to the 50% contrast static stimulus used in the main experiments (median firing rate 5.69 spikes/S, IQR 2.4-9.0 versus 4.22, IQR 2.1-7.9; $p < 0.05$, Kruskal-Wallis test). Very few units were significantly inhibited by counterphase modulated contrast increments (6.5%, 6/93 units, $p<0.05$ relative to baseline, Wilcoxon's signed rank test), whereas the vast majority of responsive units were excited (43%, 40/93 units).

The population response to contrast increments was strongly positive. However, when contrast changes were paired with optogenetic stimulation of PV neurons, the average population signal changed sign (gold). Traces depict average Gaussian filtered ($\sigma$ = 25 ms) PSTH in response to contrast increments from all units recorded with counterphase modulated in PV mice. Optogenetic stimulation produces a robust decrease in firing that is comparable in magnitude, but opposite in sign, compared to responses evoked by large contrast increments. 75 of 93 (80%) units had lower evoked responses for trials with PV stimulation and distributions of firing rate changes are significantly different (visual trials median = +1.03 ΔSpikes/s, 0.2-2.0 IQR; visual + optogenetic trials median = -0.87 ΔSpikes/s, -4.0-0.78 IQR, $p < 10^{-9}$, Komolgorov-Smirnov test). Gray square depicts the analysis window (+50 - +500 ms) used for spike rate quantification. (E) Average change in spike rate compared to the time matched baseline period for all recorded units (stimulus – baseline). The population responses were significantly different across conditions ($p < 10^{-53}$; Friedman's test). Post hoc tests indicated that the population response in each condition was significantly different from all other stimulus configurations (Dunn–Šidák correction, all comparisons $p < 0.05$). Thus, decreasing V1 spike output on the background of a dynamic visual stimulus and higher background levels of activity impairs contrast change detection. These results are consistent with our primary findings.

**Table S3**

| | 2 Hz | | | 4 Hz | | |
|---|---|---|---|---|---|---|
| **Mouse** | **Hits/Miss unstim** | **Hits/Miss stim** | **p-value** | **Hits/Miss unstim** | **Hits/Miss stim** | **p-value** |
| **1** | 146/103 | 55/170 | $< 10^{-15}$ | 273/221 | 129/371 | $< 10^{-20}$ |
| **2** | 150/120 | 70/187 | $< 10^{-11}$ | 167/236 | 83/313 | $< 10^{-10}$ |

**Table S3. Summary of changes in detection performance produced by optogenetic stimulation of PV interneurons for counterphase modulated visual stimuli.** We combined data across all sessions at the tested contrast change (+24%) for sessions in which the counterphase modulation rate was 2 Hz (left) or 4 Hz (right) and calculated the relative proportion of hits and misses separately for trials with and without optogenetic stimulation of PV interneurons. Cumulative data from the two PV mice is shown above, numbers correspond to the total counts of hits/misses for contrast increments of a 2 Hz or 4 Hz counterphase modulated stimulus. P-values correspond to the results of Fisher's exact test comparing the proportions of hits/misses with and without stimulation. Optogenetic stimulation of PV interneurons similarly impaired detection of contrast increments for both modulation rates.

**Figure S4**

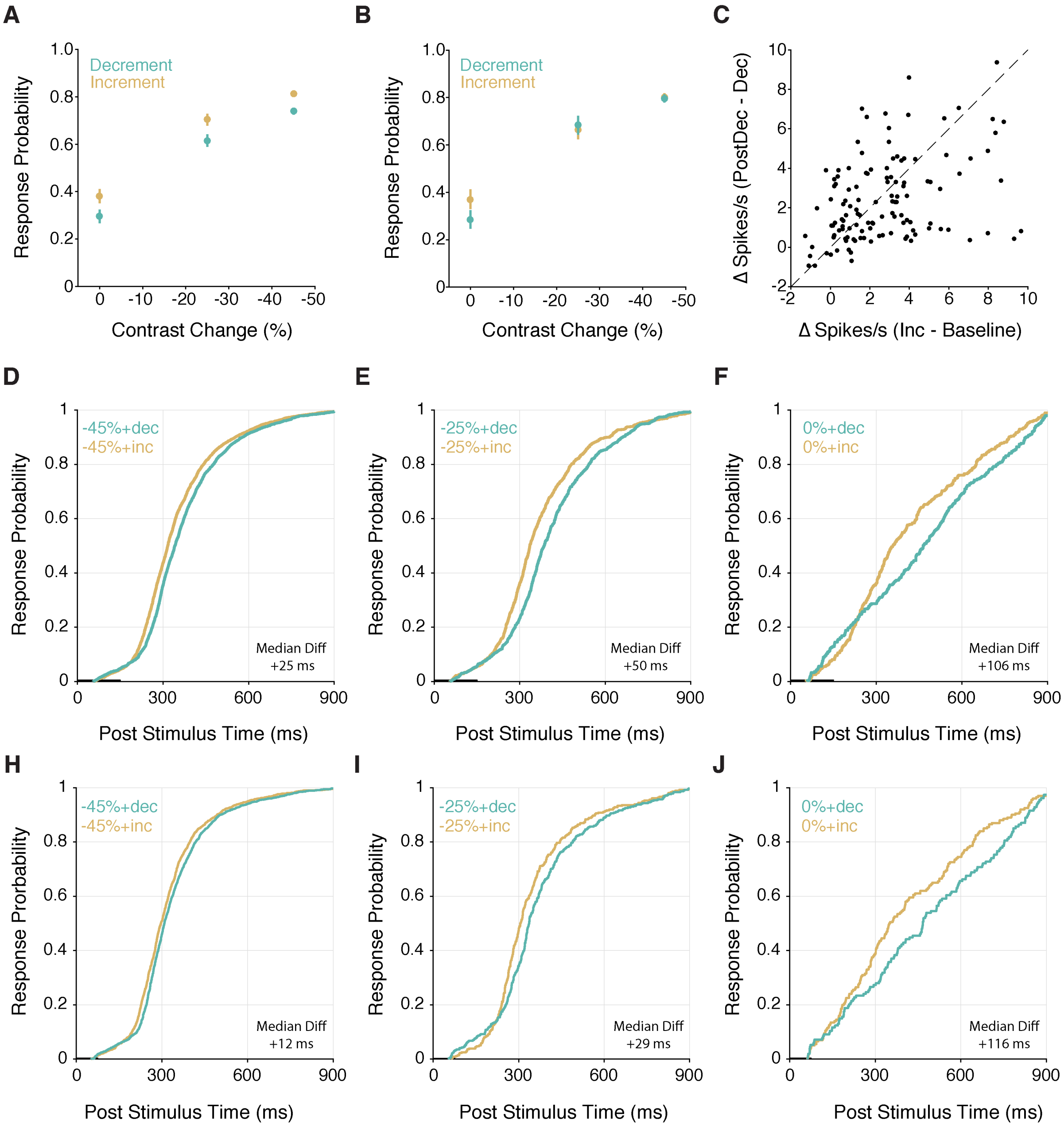


**Figure S4. Increments versus decrements in excitatory input produce dissociable effects on the probability of behavioral responses but comparable changes in spiking in V1 neurons.** In addition to trials with only changes in optogenetic stimulation, many trials featured increments and decrements in optogenetic input that were concurrently presented with large and moderate contrast decrements. We found differential effects on behavioral responses to contrast changes when those changes were paired with increases versus decreases in optogenetic input.

A) Points represent probability of a response within the 900 ms reaction time window for different visual contrast decrement magnitudes depending on whether optogenetic input decremented (aqua) or incremented (gold) for 150 ms before returning to the baseline power (n = 31 sessions in 3 mice). Mice consistently responded more when the optogenetic input incremented than when it decremented. (Response probability, -45%: increment = 0.81, 0.80-0.83 95% CI, decrement = 0.74, 0.72-0.75 95% CI; -25%: increment = 0.70, 0.68-0.73 95% CI, decrement = 0.62, 0.59-0.64 95% CI; 0%: increment = 0.38, 0.35-0.41 95% CI, decrement = 0.30, 0.27-0.32 95% CI; Effect of Contrast, $p < 10^{-20}$; Effect of Optogenetic step direction, $p < 10^{-5}$; Interaction, $p = 0.39$; logistic regression). B) Same as in A, except for optogenetic pulses of 75 ms (n = 19 sessions). Comparing across stimuli, shorter pulses had no effect on the proportion correct (Response probability, -45%: increment = 0.80, 0.78-0.82 95% CI, decrement = 0.79, 0.78-0.81 95% CI; -25%: increment = 0.66, 0.64-0.72 95% CI, decrement = 0.68, 0.64-0.72 95% CI; 0%: increment = 0.37, 0.32-0.41 95% CI, decrement = 0.28, 0.25-0.32 95% CI; Effect of Contrast, $p < 10^{-20}$; Effect of Optogenetic step direction, $p = 0.28$; Interaction, $p = 0.06$; logistic regression). However, when tested in isolation, the response probability for optogenetic increments was significantly greater than decrements when the visual stimulus did not change ($p < 0.01$; logistic regression). This shows that 75 ms optogenetic pulses were still sufficient to augment the response probability when presented in isolation.

C) We next sought to compare the magnitude of spike rate increases following optogenetic increments and cessation of optogenetic decrements (when the optogenetic input increased back to the baseline power). For each unit, we calculated the change in spike rate (Δspikes/s) during optogenetic increments relative to the prestimulus baseline (250 ms before increment onset). We also calculated the change in firing rate during the 250 ms following decrement offset relative to the decrement period. Individual points represent the change in firing rate for individual units following optogenetic increments (x-axis) and the offset of optogenetic decrements (y-axis) for equivalent step sizes (±0.1, ±0.17, ±0.28, ±0.45 mW, each unit contributes four points, one for each step size). The firing rate change following increments versus the offset of optogenetic decrements were comparable at each step size (mean Δspikes/s, [min max]; ±0.1 mW: increment = +1.11, [-1.3 +3.9], decrement offset = +0.96, [-0.9 +3.8]; ±0.17 mW: increment = +1.85, [-0.9 +4.1], decrement offset = +1.79, [-0.1 +4.5]; ±0.28 mW: increment = +3.44, [+0.2 +8.3], decrement offset = +3.14, [+0.4 +6.7]; ±0.45mW: increment = +4.3, [-3.1 +9.6], decrement offset = +3.72, [+0.4 +9.4]; all within-step size comparisons, $p > 0.05$, Wilcoxon signed-rank test). This indicates the increase in spiking compared to the immediately preceding time-matched epoch were comparable in magnitude. However, lever responses did not appear to be enhanced by the offset of the optogenetic decrement. This suggests that lever responses depending more strongly on the integrated change in spiking over longer time scales rather than instantaneous changes in V1 spiking.

D-I) Cumulative reaction time distributions on trials in which mice released the lever during the reaction time window (900 ms) following contrast changes presented concurrently with optogenetic increments and decrements. D-F) Cumulative distributions for sessions in which the optogenetic pulse duration was 150 ms when contrast changes were large (D), moderate (E), or zero (F). Lever responses were faster on trials with optogenetic increments compared to optogenetic decrements (median reaction time, -45%: increment = 317 ms, 253-413 IQR, decrement = 342, 278-439 IQR, $p < 10^{-12}$; -25%: increment = 335 ms, 266-455 IQR, decrement = 342, 308-503 IQR, $p < 10^{-9}$; 0%: increment = 358 ms, 243-583 IQR, decrement = 464, 249-665 IQR, $p < 0.01$; all Kruskal-Wallis test). Optogenetic increments presented without contrast changes (F) produce an increase in response probability compared to decrements. Thick portion of the x-axis indicates the LED pulse duration, while the visual stimulus change remained on the screen for the duration of the reaction time window (900 ms). Median Diff = difference in medians between the two reaction time distributions (increment – decrement). (G-I) Same as D-F except for sessions in which the LED pulse duration was reduced to 75 ms (19 sessions in the same 3 mice). While the distributions when visual stimuli are present are significantly different, the differences between the distributions are smaller in magnitude compared to the same stimuli presented with longer optogenetic pulse durations (median reaction times on hits, -45%: increment = 297 ms, 240-371 IQR, decrement = 309, 256-395 IQR, $p < 10^{-4}$; -25%: increment = 306 ms, 252-403 IQR, decrement = 337, 308-503 IQR, $p < 0.01$; 0%: increment = 351 ms, 220-603 IQR, decrement = 467, 276-716 IQR, $p < 0.05$; all Kruskal-Wallis test). The smaller difference between reaction time distributions is consistent with the smaller integrated difference in LED input with shorter pulses. (I) When contrast changes are absent, 75 ms optogenetic increments produce a visible increase in response probability compared to decrements.

**Figure S5**

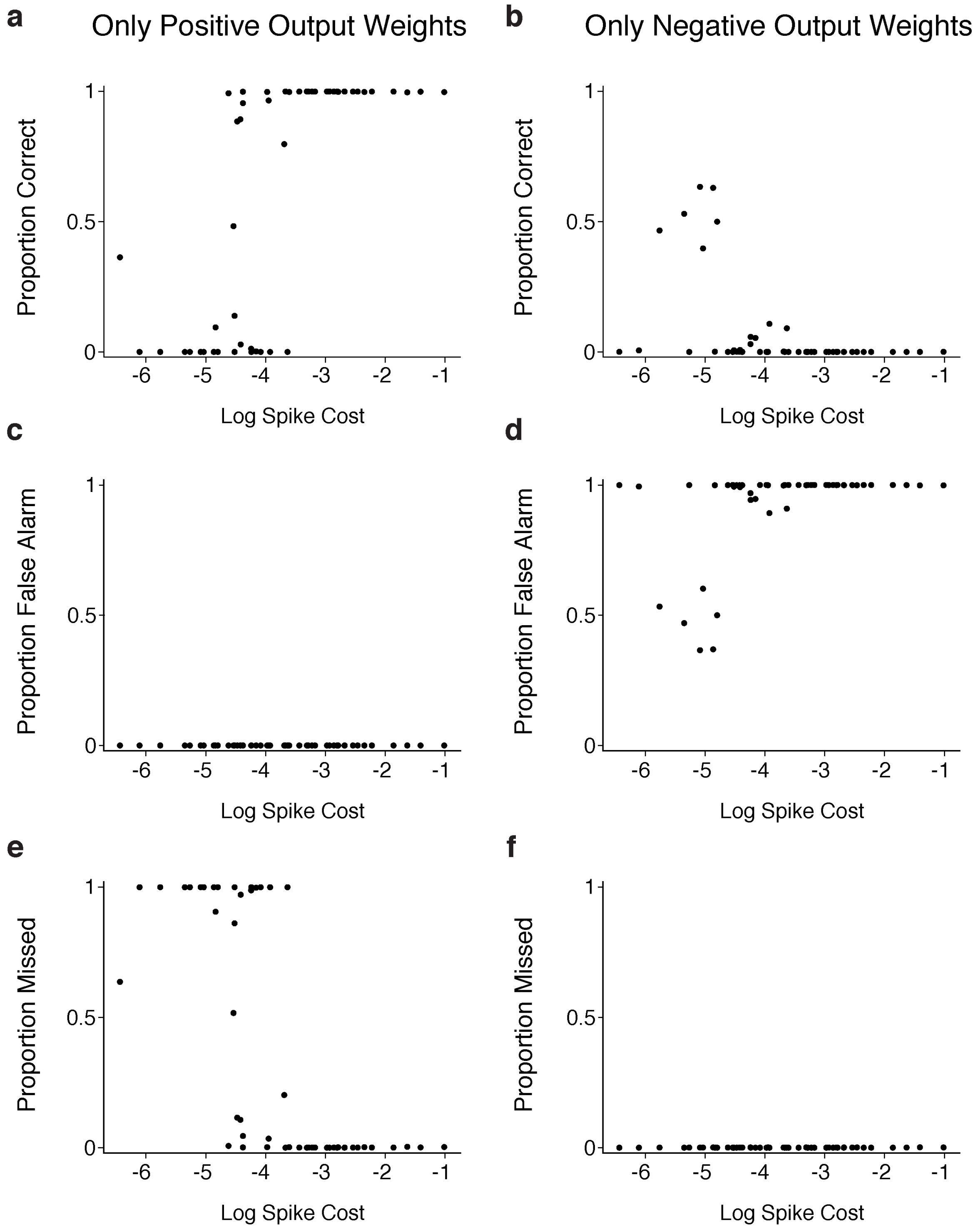
a
Only Positive Output Weights
b
Only Negative Output Weights
Proportion Correct
Proportion False Alarm
Proportion Missed
Log Spike Cost
c
d
e
f
1
0.5
0
-6
-5
-4
-3
-2
-1

**Figure S5. Contributions of positive and negative output weights to task performance in RNNs.** Following training, networks were presented with new trials and either positive or negative connections from the recurrent layer to the release neuron were shut off. (A-B) Proportion of successfully detected contrast changes for as a function of spike cost when trained networks must perform the task using exclusively positive (A) or negative (B) output weights from the recurrent layer. Each point represents the performance of a single trained network. RNNs perform well using positive but not negative output weights across a range of spike costs. (C-D) Proportion of false alarms as a function of spike cost when RNNs use only positive (C) or negative (D) output weights. Networks restricted to exclusively negative weights exhibit high false alarm rates. (E-F) Proportion of contrast changes that networks failed to detect (misses) as a function of spike cost for RNNs restricted to exclusively positive (E) or negative (F) output weights.
